# A structure-selective endonuclease drives uniparental mitochondrial DNA inheritance

**DOI:** 10.64898/2026.03.19.713005

**Authors:** Mayu Shimomura, Hwa Young Yun, Philip C. Zuzarte, Jared T. Simpson, Haley D. M. Wyatt, Thomas R. Hurd

**Affiliations:** Department of Molecular Genetics, University of Toronto; 661 University Avenue, Toronto, Ontario, M5G 1M1, Canada; Department of Biochemistry, University of Toronto; 661 University Avenue, Toronto, Ontario, M5G 1M1, Canada; Ontario Institute for Cancer Research; 661 University Avenue, Toronto, Ontario, M5G 1M1, Canada; Genetics and Genome Biology Program, Hospital for Sick Children; 686 Bay St., Toronto, Ontario, M5G 0A4, Canada

**Keywords:** Mitochondria, mitochondrial DNA (mtDNA), maternal mtDNA inheritance, paternal mtDNA elimination, testis, spermatogenesis, endonuclease, GIY-YIG endonuclease

## Abstract

Maternal inheritance of mitochondrial DNA (mtDNA) is a near-universal feature of eukaryotes^1^, yet the mechanisms that ensure this by preventing paternal mtDNA inheritance have remained unclear. In both *Drosophila* and humans, mtDNA is actively eliminated from sperm during spermatogenesis, producing mature sperm whose mitochondria lack their genomes^2–5^. Here we identify Hotaru, a previously uncharacterized, testis-specific GIY-YIG endonuclease, as a central player in this process. We find that Hotaru is expressed in elongated spermatids, localizes to the mitochondrial matrix, and is required for paternal mtDNA elimination. In *hotaru* mutants, sperm retain mtDNA at levels comparable to those present before the elimination process. Genetic and biochemical analyses show that Hotaru selectively recognizes and cleaves cruciform DNA structures within the mtDNA control region. Together, these findings identify a dedicated nuclease that enforces mitochondrial genome elimination in the animal male germline and reveal that an unexpected structural feature of mtDNA serves as the molecular determinant of its destruction. By recognizing DNA structure rather than specific sequence motifs, this mechanism is inherently robust to the high mutation rate of mitochondrial genomes.

---

Mitochondria are unique among animal organelles in that they retain their own genomes, a vestige of their proteobacterial origins^6,7^. In most animals, including humans, mtDNA exists as a small, circular molecule that encodes core components of the oxidative phosphorylation machinery, required for ATP production^8,9^. mtDNA is packaged with the DNA-binding protein TFAM into a structure known as a nucleoid^10^. Unlike nuclear DNA, mtDNA is present in hundreds to thousands of copies in most cells^11^, and its replication is not tightly coupled to the cell cycle^12,13^. Given these fundamental differences, it is not surprising that mtDNA follows a distinct inheritance pattern from nuclear DNA. During sexual reproduction, offspring inherit nuclear genomes from both parents, but the mitochondrial genome is inherited exclusively through the egg. Despite the near universality of maternal inheritance, the evolutionary forces that favour it, and the cellular mechanisms that eliminate paternal mtDNA, remain largely unresolved.

Many organisms, including *Drosophila*, mice, and humans, eliminate paternal mtDNA during spermatogenesis while retaining the mitochondria themselves^2,3,14,15^. Recent work has linked this removal to a marked reduction in TFAM levels during spermatogenesis in both flies and mammals^2,15–17^. However, the active mechanisms that degrade mtDNA remain unclear. Two proteins have previously been implicated in the nucleolytic degradation of mtDNA. In *D. melanogaster* and *C. elegans*, the intermembrane-space endonuclease EndoG affects the timing of mtDNA removal^5,18^. However, it resides in a different mitochondrial compartment from mtDNA^19,20^ (Extended Data Fig. 1B), and its loss only delays rather than prevents mtDNA elimination^5,14^. In *D. melanogaster*, POLDIP2, a substrate-specific adaptor of the ClpXP protease^21^, has also been recently implicated in eliminating paternal mtDNA. Although one study proposed that POLDIP2 possesses exonuclease activity, POLDIP2 lacks recognizable catalytic motifs characteristic of known exonucleases^3^. Another study showed that POLDIP2 functions as a ClpXP substrate adaptor required for TFAM downregulation^15^, a step that would increase mtDNA accessibility and thereby facilitate its elimination^22^. Together, these findings suggest that the key nuclease responsible for directly cleaving and degrading paternal mtDNA has yet to be identified.

## Identification of Hotaru, a mitochondrial matrix-localized endonuclease

Given that mtDNA is circular and resides within the mitochondrial matrix, we hypothesized that a matrix-localized endonuclease must first trigger its elimination. We therefore screened Drosophila endonucleases for matrix localization using a split-GFP assay^23^. We tethered a GFP fragment (GFP1-10) to TFAM, which localizes to the matrix, and the complementary fragment (GFP11, in seven tandem repeats) to candidate endonucleases. Endonucleases that localize to the matrix bring the two GFP fragments into proximity, resulting in GFP reconstitution and green fluorescence in co-expressing cells (Fig. 1A, cartoon). Of the 27 endonucleases screened, an uncharacterized endonuclease CG42391, which we named Hotaru (see below), produced the highest proportion of GFP-positive cells, followed by Fen1 and Ogg1, both of which have previously been shown to localize to mitochondria^24,25^ (Fig. 1A, B, and Extended Data Fig. 1A). In contrast, EndoG, which has been implicated in paternal mtDNA elimination^5^, did not show detectable matrix localization (Fig. 1A). Only when we expressed it with GFP1-10 fused to the C-terminus of Opa1, a known intermembrane space protein, did we observe clear GFP fluorescence (Extended Data Fig. 1B), consistent with intermembrane space localization.

**Fig. 1.**
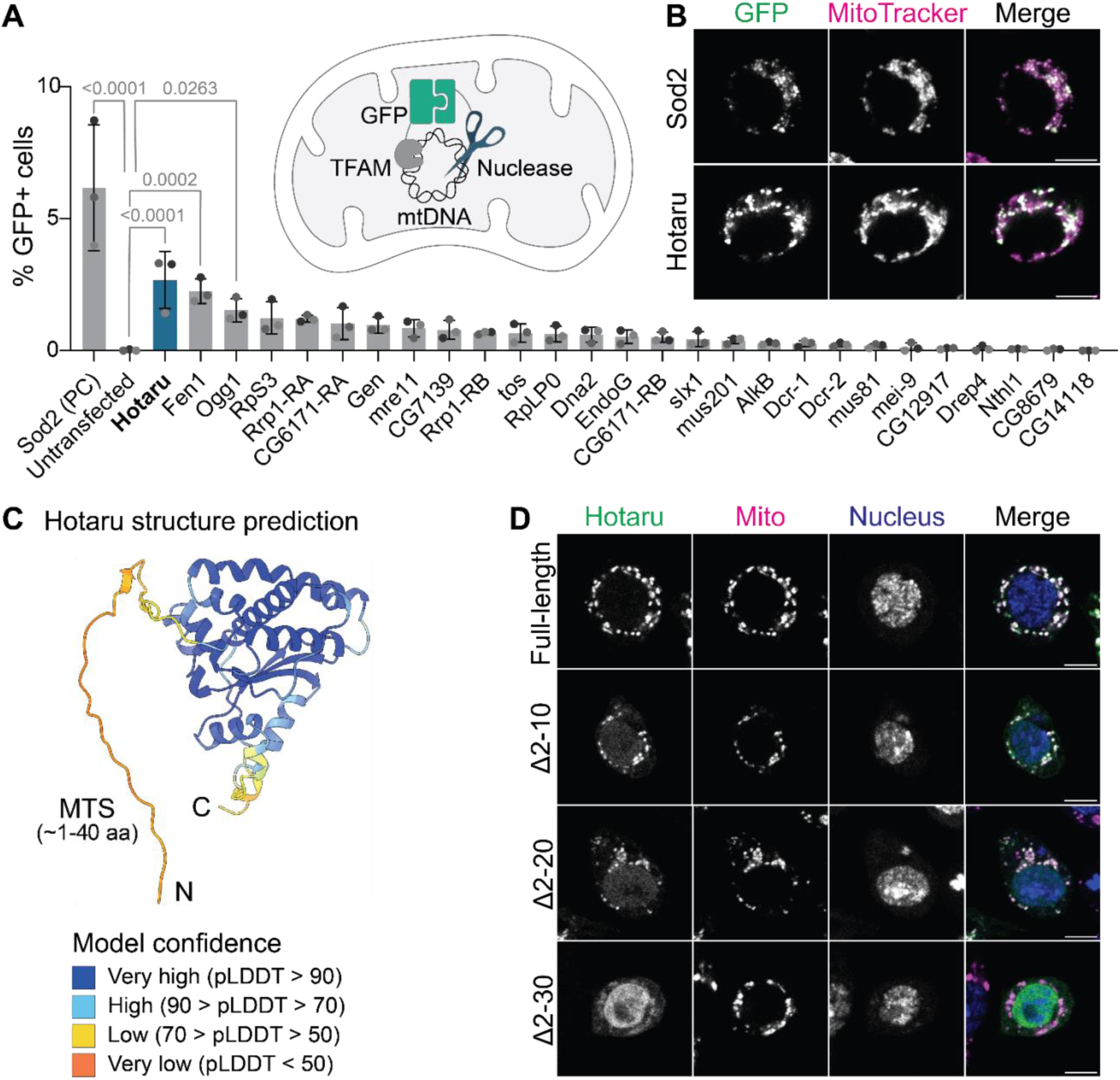
| Split-GFP screen identifies Hotaru as a mitochondrial endonuclease. (**A**) Schematic of the split-GFP assay used to assess nuclease localization. GFP1–10 and GFP11 reconstitute a fluorescent chromophore only when the candidate nuclease localizes to the mitochondrial matrix. Bar graph shows the percentage of GFP-positive cells co-transfected with TFAM::GFP1–10 and the indicated nuclease fused to seven tandem GFP11 tags. PC, positive control. n = 3 independent transfections. One-way ANOVA with Dunnett’s test. Data, mean ± s.d. (**B**) Confocal micrographs of S2 cells expressing TFAM::GFP1–10 with either Sod2::GFP11×7 (positive control) or Hotaru::GFP11×7. Reconstituted GFP signal colocalizes with mitochondria labelled with MitoTracker Red CMXRos. Max intensity projections of three slices across 0.595 μm. Scale bars, 5 μm. (**C**) AlphaFold prediction of full-length Hotaru (AF-BZYZH2-F1) coloured by per-residue confidence score (pLDDT). The N-terminal ∼40 amino acids are predicted to contain a mitochondrial targeting sequence (MTS). (**D**) Confocal micrographs of S2 cells expressing N-terminally truncated Hotaru. Full-length Hotaru robustly colocalizes with mitochondria marked with anti-ATP5A, whereas progressive N-terminal deletions decrease this colocalization. Nucleus is marked with DAPI. Scale bars, 5 μm.

Computational analysis predicted Hotaru to possess an N-terminal mitochondrial targeting sequence (MTS) within approximately the first 40 amino acids (probability = 0.8998, MitoProt II v.1.101^26^; Fig. 1C). Indeed, N-terminal truncations markedly impaired Hotaru’s targeting to mitochondria as measured using both the split-GFP assay (Extended Data Fig. 1C) and immunofluorescence (Fig. 1D). Together, these findings identify Hotaru as a mitochondrial matrix-localized endonuclease and a candidate for initiating paternal mtDNA elimination.

## Hotaru is required for paternal mtDNA elimination

To investigate *hotaru*’s relevance for paternal mtDNA elimination, we stained mature sperm from controls and various *hotaru* mutants with the DNA intercalating dye DAPI. We first analyzed *hotaru* mutants carrying a premature stop codon caused by a gene trap insertion^27^ *in trans* to a large deletion that removes the entire gene locus and adjacent regions (Fig. 2A). In control sperm, DAPI staining was restricted to needle-shaped nuclear DNA, consistent with complete elimination of paternal mtDNA^5,14^ (Fig. 2B). In contrast, these *hotaru* mutants displayed distinct DAPI-positive foci along the sperm tail, through which the mitochondrial derivatives run, suggesting that paternal mtDNA elimination was defective (Fig. 2B, D). To validate this phenotype, we generated *hotaru* knockout (KO) flies using CRISPR-Cas9 gene editing (Fig. 2A); these flies similarly exhibited a significant increase in non-nuclear DAPI foci (Fig. 2C, E). We did not observe DAPI-positive foci in germline knockdowns of the other two mitochondrial matrix-localized endonucleases identified in our split-GFP assay (Fen1 and Ogg1) nor in *endoG* mutants^5^ (Extended Data Fig. 2). Together, these data indicate that *hotaru* is required for paternal mtDNA elimination. We named CG42391 “Hotaru”, the Japanese word for “firefly”, because the persisting punctate DAPI foci in the mutants resemble fireflies glowing in the night.

**Fig. 2.**
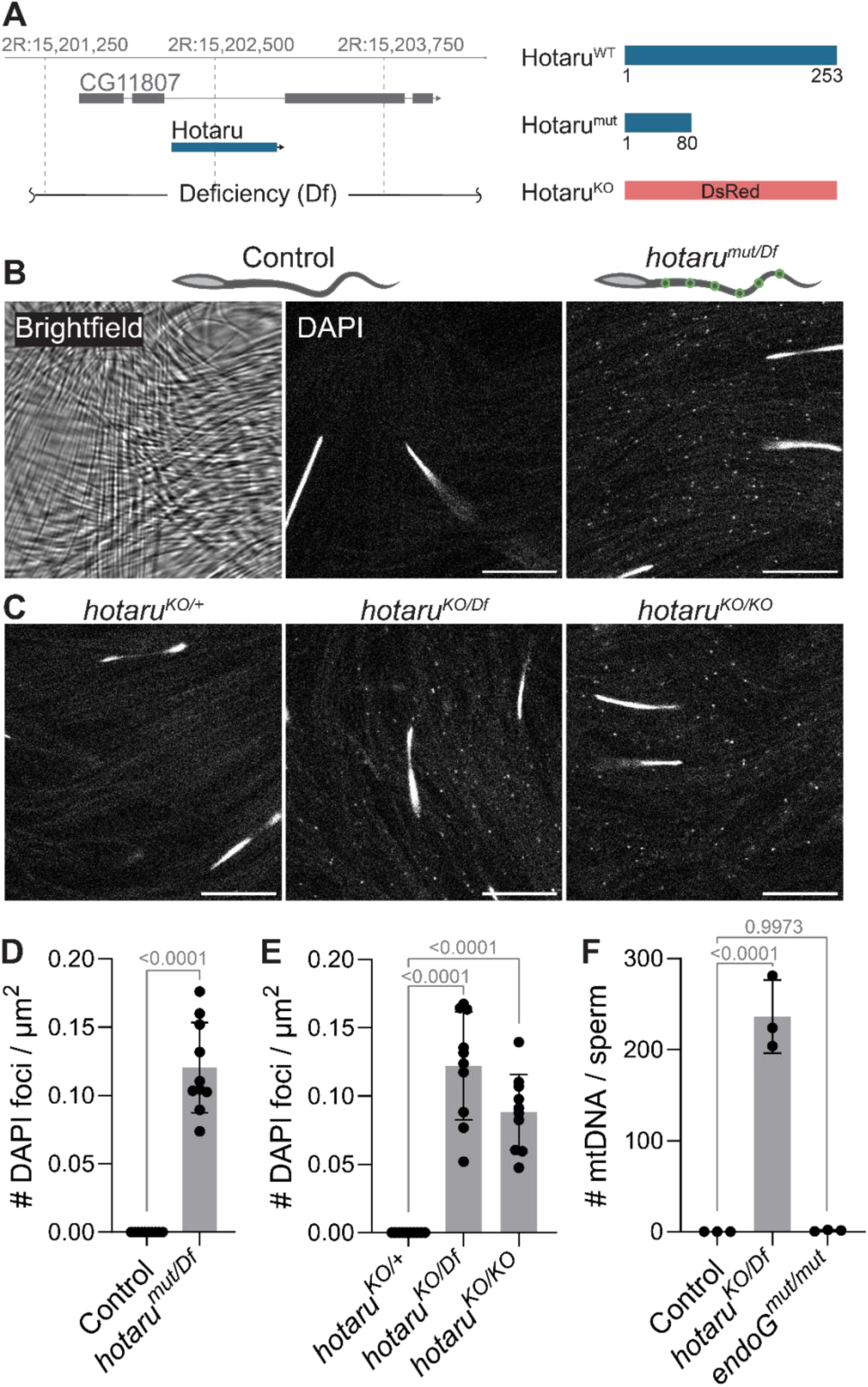
| Hotaru is required for paternal mtDNA elimination. (**A**) Genomic organization of the *hotaru* locus and schematic representation of alleles used in this study. *hotaru^mut^*carries a CRIMIC insertion introducing a premature stop codon upstream of the catalytic GIY-YIG endonuclease domain. *hotaru^KO^* replaces the entire coding sequence with dsRed. *hotaru^Df^* removes a genomic region (∼133 kb) including the entire *hotaru* locus. (**B**) Confocal micrographs of mature sperm from control (*w^1118^*) (middle) and *hotaru^mut/Df^* transheterozygote (right) stained with DAPI, which marks both the needle-shaped nuclear DNA and uneliminated mtDNA. A brightfield image of control sperm is also included (left). Scale bars, 10 μm. (**C**) Confocal micrographs of DAPI-stained sperm from heterozygous and homozygous *hotaru^KO^* males. Scale bars, 10 μm. (**D**) Quantification of DAPI-positive mtDNA foci for the genotypes shown in (B). n = 10 independent males. Two-tailed Student’s t-test. Data, mean ± s.d. (**E**) Quantification of DAPI-positive mtDNA foci for the genotypes shown in (C). n = 10 independent males. One-way ANOVA with Dunnett’s test. Data, mean ± s.d. (**F**) mtDNA copy number per sperm measured by ddPCR. Each data point represents sperm transferred to 20 females from 15 males. One-way ANOVA with Dunnett’s test. Data, mean ± s.d.

To quantify how much mtDNA remains when Hotaru is eliminated, we assessed mtDNA copy number in mature sperm from control and *hotaru* KO males using droplet digital PCR (ddPCR) (Extended Data Fig. 3). In controls and *endoG* mutants, paternal mtDNA levels were zero, as expected (Fig. 2F). In contrast, *hotaru* KO sperm contained abundant mtDNA, with each containing approximately 240 mtDNA copies (Fig. 2F). This number closely matches the total mtDNA content prior to the elimination phase^5,28^. These results indicate that most, if not all, mtDNA is retained in mature sperm when *hotaru* function is lost.

## Hotaru acts in elongated spermatids to eliminate paternal mtDNA

If Hotaru drives the paternal mtDNA elimination, it must be expressed in the testis and act no later than the stages when mtDNA is removed. To determine whether Hotaru is expressed in the appropriate tissue at the appropriate time, we first assessed *hotaru* mRNA expression. RNA-sequencing data revealed strong testis enrichment^29^ (Fig. 3A), and RNA fluorescent *in situ* hybridization (FISH) detected *hotaru* mRNA in the germline throughout spermatogenesis (Fig. 3B, C, Extended Data Fig. 4).

**Fig. 3.**
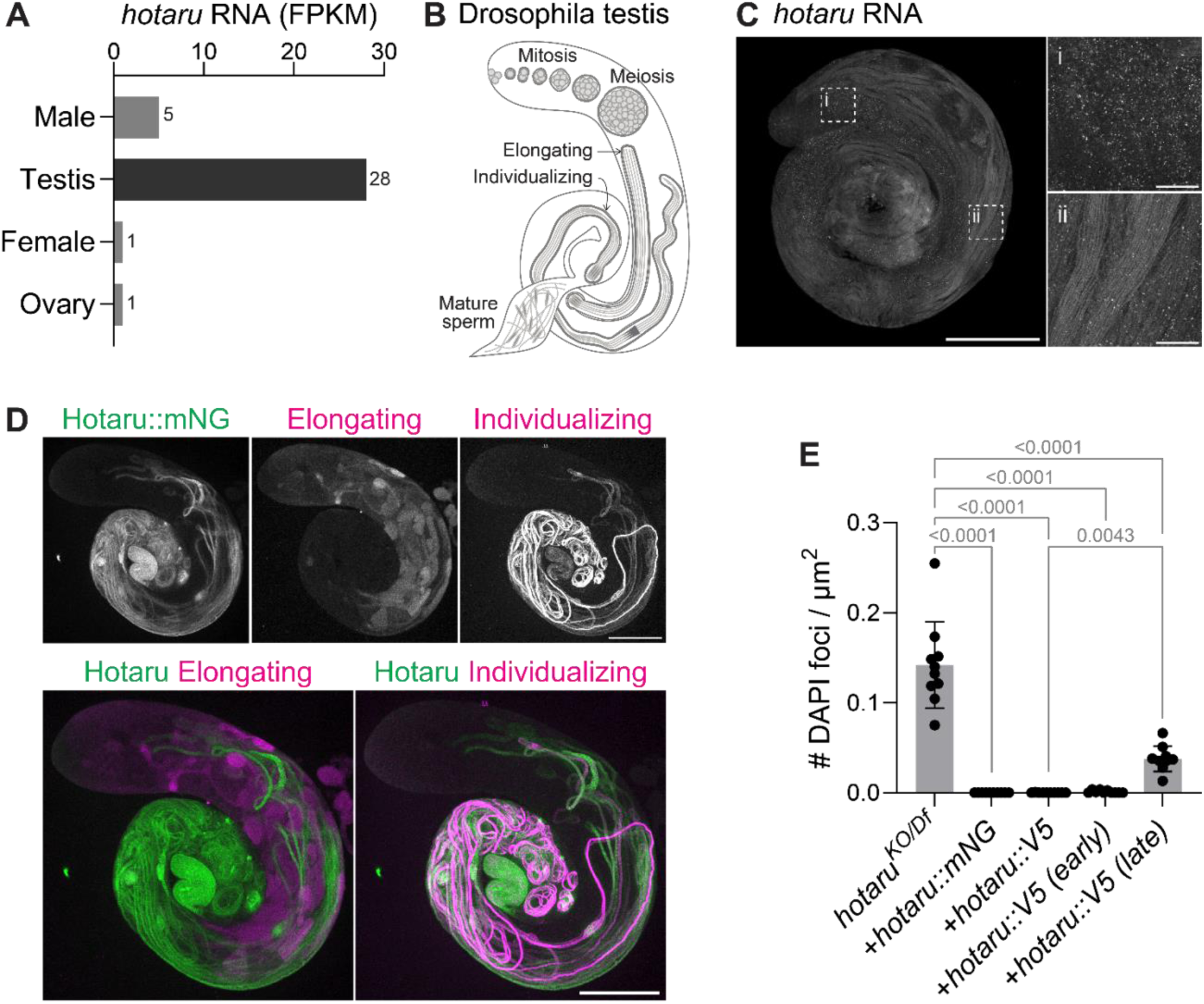
| Hotaru functions in elongated spermatids. (**A**) Expression of *hotaru* RNA across sex and tissues based on FlyAtlas2 RNA-seq data^29^. (**B**) Schematic of Drosophila spermatogenesis. Germline stem cells (GSCs) produce gonialblasts, which undergo mitotic and meiotic divisions to generate cysts of 64 spermatids. These spermatids elongate, and once fully elongated, the spermatids are separated through the process of individualization to form mature sperm. (**C**) HCR-FISH detection of *hotaru* mRNA in control (*w^1118^*) testes. *hotaru* transcripts are present throughout spermatogenesis including in spermatocytes (i) and elongating/elongated spermatids (ii). Scale bars: whole testis, 200 μm; insets, 20 μm. (**D**) Confocal micrographs of Hotaru::mNeonGreen (mNG) co-stained with the late-elongating cyst marker anti-Soti and the individualizing cyst marker anti-polyglycylated tubulin. Polyglycylated tubulin is present from the onset of individualization to mature sperm. Max intensity projections across the entire thickness of testes. Scale bars, 200 μm. (**E**) Quantification of DAPI-positive mtDNA foci for the rescue constructs. *hotaru::V5 (early)* and *hotaru::V5 (late)* is expressed under the 5’ UTR of *CG42355* and *mst87F*, respectively. n = 10 independent males. One-way ANOVA with Tukey’s test. Data, mean ± s.d. Representative images are shown in Extended Data Fig. 5B.

We next examined Hotaru protein expression, as transcript abundance poorly predicts protein levels in the Drosophila testis, especially in post-meiotic spermatids, which are transcriptionally silent and extensively post-transcriptionally regulated^30^. Previous work indicates that paternal mtDNA elimination occurs just before the individualization of spermatids to generate mature sperm^5,14^. To determine when Hotaru protein appears, we tagged Hotaru with mNeonGreen and expressed it under its own regulatory sequences (Extended Data Fig. 5A). This transgene fully rescued mtDNA elimination in *hotaru* KO males, confirming the tagged protein’s functionality (Fig. 3E, Extended Data Fig. 5B). Co-staining showed that Hotaru protein colocalizes with individualizing cyst marker polyglycylated tubulin^31^ but not the late elongating cyst marker Soti^31^, indicating that it is first detectable in elongated spermatids at the onset of individualization (Fig. 3D, Extended Data Fig. 4B). This expression pattern coincides exactly with the developmental window during which paternal mtDNA is eliminated.

We next asked whether Hotaru is required at this stage. We performed rescue experiments by reintroducing Hotaru into *hotaru* KO males using regulatory elements of two post-meiotically translated genes^32,33^. The *CG42355* regulatory elements drove expression in early spermatids prior to elongation and the *mst87F* regulatory elements drove expression only after individualization has begun (Extended Data Fig. 5C). Expression of *hotaru* under its native regulatory elements or under those of *CG42355* fully restored paternal mtDNA elimination (Fig. 3E, Extended Data Fig. 5B). In contrast, expression under *mst87F* UTR failed to fully rescue the defect (Fig. 3E, Extended Data Fig. 5B). These results demonstrate that Hotaru is required in spermatids by the onset of individualization, when paternal mtDNA elimination occurs.

## Catalytic activity of Hotaru is essential for mtDNA cleavage and elimination

Hotaru is predicted to contain a GIY-YIG endonuclease domain^34,35^, comprising a three-stranded antiparallel β-sheet flanked by α-helices^36^ (Extended Data Fig. 6). The first two β-strands contain the conserved catalytic tyrosines of the GIY and YIG motifs (Y88 and Y121 in Hotaru; Figure 4A, Extended Data Fig. 6). Additional conserved active-site residues include an invariant Glu^36^ (Extended Data Fig. 6), corresponding to E186 in Hotaru (Fig. 4A). To date, two animal GIY-YIG endonucleases, SLX1 and ANKLE1, have been biochemically characterized, both of which display DNA structure-selective endonuclease activity^37–42^. To assess whether Hotaru exhibits similar substrate selectivity, we examined its activity *in vitro*. We purified full-length (FL) Hotaru, the mature form lacking its MTS (ΔMTS; Δ2-41), and a catalytically-dead Y88F mutant from baculovirus-infected insect cells (Extended Data Fig. 7) and assayed these against a panel of linear and branched DNA substrates (Extended Data Fig. 8)^37,38,43^. Wild-type Hotaru (FL and ΔMTS) cleaved three-way junctions (splayed arm, 5’-flap, and 3’-flap) and a four-way nicked Holliday junction but showed no activity on linear DNA (Fig. 4B, Extended Data Fig. 8). This substrate preference resembles that of SLX1 and ANKLE1^37–42^. Notably, however, Hotaru did not cleave an intact Holliday junction (Extended Data Fig. 8), revealing a substrate selectivity distinct from these enzymes. As expected, the Y88F mutant lacked detectable nuclease activity on all substrates tested (Fig. 4B, Extended Data Fig. 8).

**Fig. 4.**
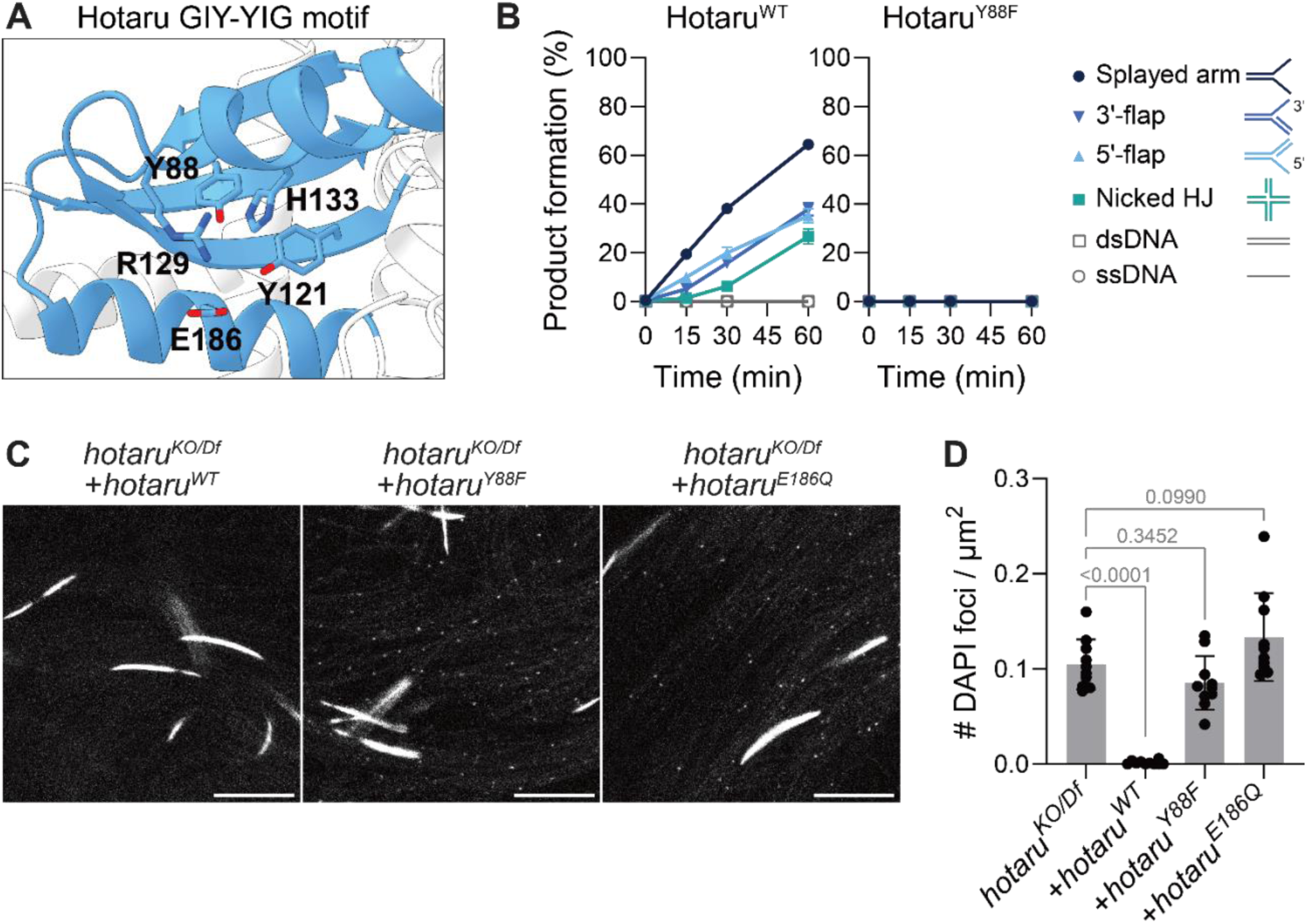
| Hotaru cleaves and eliminates mtDNA via its endonuclease activity. (**A**) The conserved active site residues in the predicted GIY-YIG nuclease domain of Hotaru (AF-BZYZH2-F1) spanning residues 85-244 (Pfam PF22945)^34,35^. Key residues are labelled and shown in stick representation to illustrate their spatial arrangement. (**B**) Quantification of *in vitro* nuclease activity of wild-type Hotaru (ΔMTS) and catalytically-dead Hotaru Y88F on various DNA substrates (Extended Data Fig. 8). Product formation is expressed as a percentage of total DNA. n = 3 independent assays. Data, mean ± s.d. (**C**) Confocal micrographs of DAPI-stained sperm from *hotaru^KO/Df^* males expressing *UAS-hotaru* (*WT, Y88F,* or *E186Q*) in the germline using *bam-GAL4*. Scale bars, 10 μm. (**D**) Quantification of DAPI-positive mtDNA foci for the genotypes shown in (C). n = 10 independent males. One-way ANOVA with Dunnett’s test. Data, mean ± s.d.

Having identified key catalytic residues, we next tested whether Hotaru’s enzymatic activity is required for paternal mtDNA elimination *in vivo*. Germline expression of wild-type Hotaru fully rescued mtDNA elimination in *hotaru* KO males (Fig. 4C, D). In contrast, the catalytically-dead Y88F and E186Q variants failed to do so (Fig. 4C, D), despite normal protein levels and mitochondrial localization (Extended Data Fig. 9A, B). Moreover, expression of wild-type Hotaru during early spermatogenesis, prior to its normal expression window, significantly reduced mtDNA copy number in the testis, whereas catalytically-dead mutants had no effect (Extended Data Fig. 9C). Together, these findings demonstrate that Hotaru functions as a mitochondrial endonuclease whose catalytic activity is required for paternal mtDNA elimination, consistent with a model in which Hotaru directly cleaves paternal mtDNA to initiate its destruction.

## Ectopic Hotaru expression induces mtDNA destruction

We next asked whether Hotaru is sufficient to drive mtDNA destruction in other tissues. To test this, we ectopically expressed Hotaru in the ovary, the tissue with the highest mtDNA abundance^44^. Strikingly, expression of Hotaru in female germline stem cells from the beginning of oogenesis caused severe defects in germline development, precluding analysis of mtDNA copy number, and this phenotype depended on its catalytic activity (Extended Data Fig. 9D). By contrast, restricting Hotaru expression to later stages of oogenesis largely preserved germline development. Under these conditions, ovarian mtDNA copy number was significantly reduced (Fig. 5A). This reduction again depended on Hotaru’s catalytic activity, as the catalytically-dead variants did not reduce mtDNA levels (Fig. 5A). These results demonstrate that Hotaru’s endonuclease activity is sufficient to drive mitochondrial genome elimination when ectopically expressed.

**Fig. 5.**
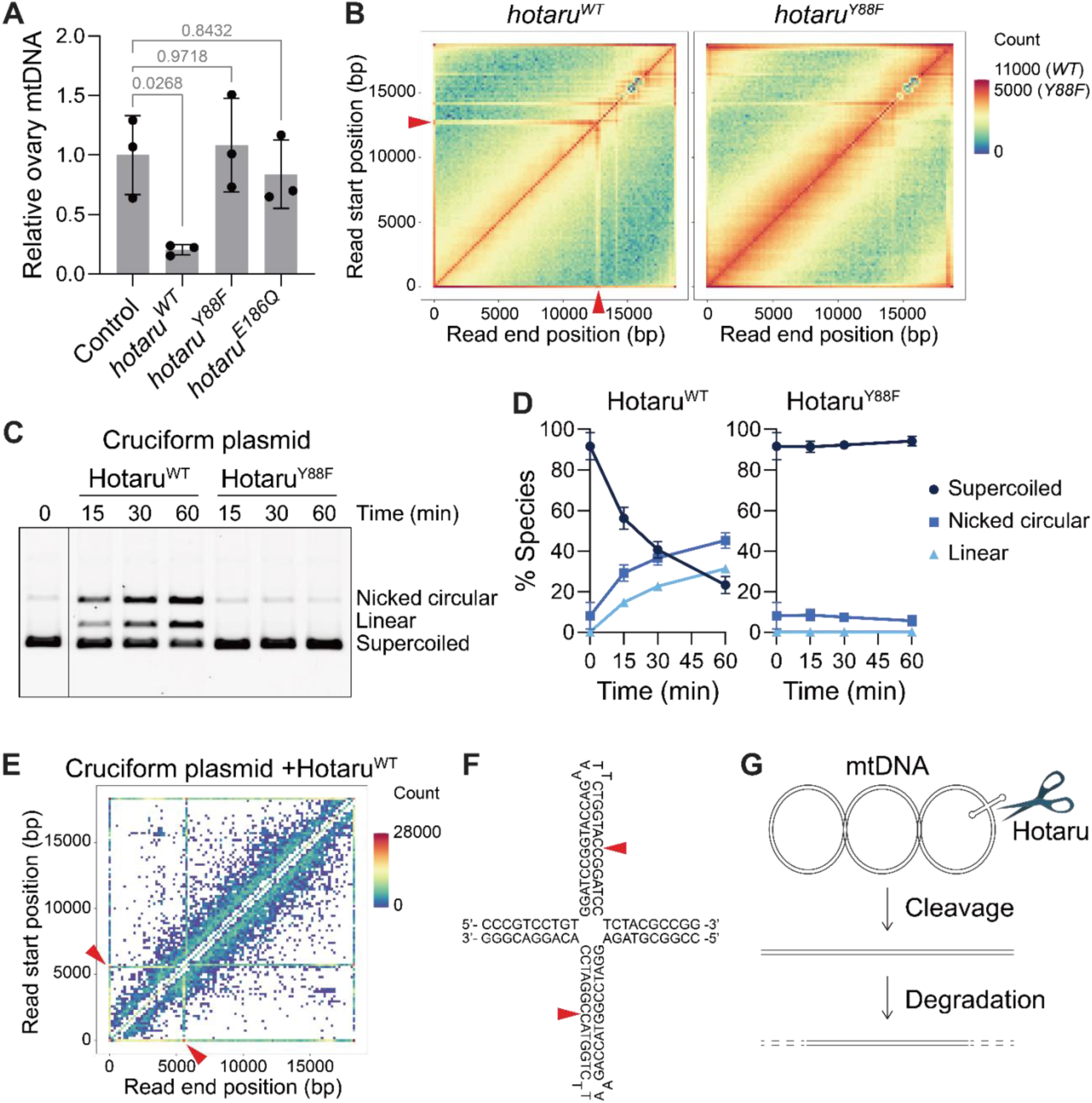
| Hotaru cleaves cruciform structures in the control region of mtDNA. (**A**) mtDNA copy number in ovaries expressing control (*UAS-mito::mCherry*) or *UAS-hotaru* (*WT, Y88F,* or *E186Q*) using *matα-tub-GAL4*. mtDNA levels were measured by qPCR using mitochondrial (COX1, COX3) primers and normalized to nuclear genes (RpL11, tub). Each data point represents a pool of three pairs of ovaries. One-way ANOVA with Dunnett’s test. Data, mean ± s.d. (**B**) Heat map generated from ONT reads of mtDNA isolated from eggs expressing *UAS-hotaru* (*WT or Y88F*) using *matα-tub-GAL4*. A prominent cleavage hotspot (arrowheads) is detected in the control region of mtDNA in wild-type samples. (**C**) Cruciform plasmid (pIRbke8^mut^)^45–47^ incubated with purified wild-type Hotaru (ΔMTS) or catalytically-dead Hotaru Y88F for 0 (no enzyme), 15, 30, and 60 min. (**D**) Quantification of (C). n = 4 independent assays. Data, mean ± s.d. (**E**) Heat map generated from ONT reads of cruciform plasmid after 60 min incubation with Hotaru. Prominent cleavage hotspots (arrowheads) are detected in the cruciform-forming inverted repeat. (**F**) Cleavage site structure modeled from read start/end site sequences. Hotaru is predicted to introduce symmetric cuts within the cruciform stem (arrowheads). (**G**) Model of mtDNA elimination during Drosophila spermatogenesis.

## Hotaru cleaves cruciform structures in the mtDNA control region

We sought to identify how Hotaru recognizes and cuts mtDNA. To map Hotaru cleavage sites, we developed a long-read Oxford Nanopore Technologies (ONT) sequencing approach. We isolated mtDNA from samples expressing either wild-type or catalytically-dead Hotaru and performed ONT sequencing, reasoning that Hotaru cleavage sites would be marked by reads initiating or terminating in the wild-type samples but not in catalytically-dead controls.

As it is not possible to recover sufficient mtDNA from the male germline, we instead analyzed mtDNA from eggs from ovaries expressing either wild-type or catalytically-dead Hotaru. Catalytically-dead samples showed no prominent cleavage sites whereas wild-type samples exhibited a prominent cleavage hotspot (Fig. 5B). This site mapped to the control region of the mitochondrial genome, rich in inverted repeats (Extended Data Fig. 10A). We detected reads initiating and terminating on both DNA strands, suggesting that Hotaru generates double-strand breaks in this region. Given the propensity of inverted repeats to form cruciform structures, together with Hotaru’s substrate selectivity toward structured DNA, we hypothesized that Hotaru targets cruciforms. To test this, we assessed its activity on a model plasmid containing an inverted repeat of a sequence distinct from that in mtDNA but capable of forming a cruciform^45–47^ In the absence of Hotaru, the plasmid migrated predominantly as a supercoiled species, indicating no detectable cleavage. In contrast, incubation with Hotaru resulted in a marked accumulation of nicked open-circular and linear species, consistent with efficient nicking of one or both DNA strands, respectively (Fig. 5C, D, Extended Data Fig. 10B, C). ONT sequencing of cleavage products confirmed that cleavage occurred in the cruciform itself, revealing that Hotaru generates symmetric cuts within the cruciform stem (Fig. 5E, F, Extended Data Fig. 10D), which we predict produces fragments with 3’ overhangs that are removed during sequencing library preparation to create the observed reads. In sum, these findings support a model in which Hotaru recognizes cruciform structures in the mtDNA control region and introduces breaks that prime the mitochondrial genome for downstream degradation (Fig. 5G).

## DISCUSSION

Reduction of mtDNA copy number during spermatogenesis is widely conserved across animals, although how this occurs has remained poorly understood^48^. Here we identify Hotaru, a mitochondrial endonuclease that directly cuts mtDNA in the male germline of *Drosophila*, establishing endonuclease-mediated programmed DNA breakage as the key initiating step in paternal mtDNA elimination. Unexpectedly, Hotaru recognizes mtDNA by targeting cruciform structures within the control region, demonstrating that the topology of the mitochondrial genome intrinsically encodes a cue for its own destruction. This reveals a previously unrecognized role for higher-order genome architecture as a determinant of genetic transmission. By targeting DNA structures rather than specific sequence motifs, this mechanism can confer robustness, as single nucleotide mutations would be less likely to disrupt cleavage.

An important question moving forward is how Hotaru is integrated into the broader machinery that enforces paternal mtDNA elimination. Cruciform extrusion within the control region likely depends on negative supercoiling of mtDNA, a topological state promoted by TFAM binding^49^. This raises the possibility that TFAM not only compacts the mitochondrial genome but also licenses its destruction by facilitating the formation of DNA structures recognized by Hotaru. In such a model, dynamic regulation of TFAM could coordinate a two-step process: structural priming of cleavage through topology modulation, followed by TFAM depletion to expose DNA for downstream degradation. Indeed, recent studies linking TFAM depletion to paternal mtDNA elimination in both flies and mammals^2,15–17^ implicate nucleoid organization as a regulatory switch governing mitochondrial genome fate. What occurs after Hotaru-mediated incision remains undefined, but the generation of DNA ends would render the mtDNA susceptible to progressive clearance by exonucleases. Together, these findings support a model in which mtDNA topology and nucleoid organization are not merely structural attributes, but active determinants of selective genome elimination.

Equally important will be understanding the fitness consequences of impairing paternal mtDNA elimination. The near-universal reduction of paternal mtDNA during spermatogenesis across animals suggests strong selective pressure for this process. Although a failure to remove mtDNA from sperm has been associated with infertility in flies and mammals^3,50^, males lacking *hotaru* exhibited no detectable reduction in fertility (Extended Data Fig. 11C). This finding suggests that the evolutionary significance of paternal mtDNA elimination may lie beyond acute effects on reproduction. Notably, the availability of *hotaru* mutants now provides an opportunity to directly test the longer-term consequences of paternal mtDNA elimination failure at the organismal and population levels.

Our findings suggest that nuclease-mediated genome destruction represent a conserved principle of paternal mitochondrial genome elimination across animals, including mammals. Because mammalian mtDNA is similarly eliminated during spermatogenesis and shares the same circular topology, its destruction is likely also initiated by an endonuclease. Recognition of DNA structure rather than sequence, as observed for Hotaru, would provide a robust strategy for targeting rapidly evolving, highly polymorphic mitochondrial genomes.

---

**Extended Data Fig. 1.**
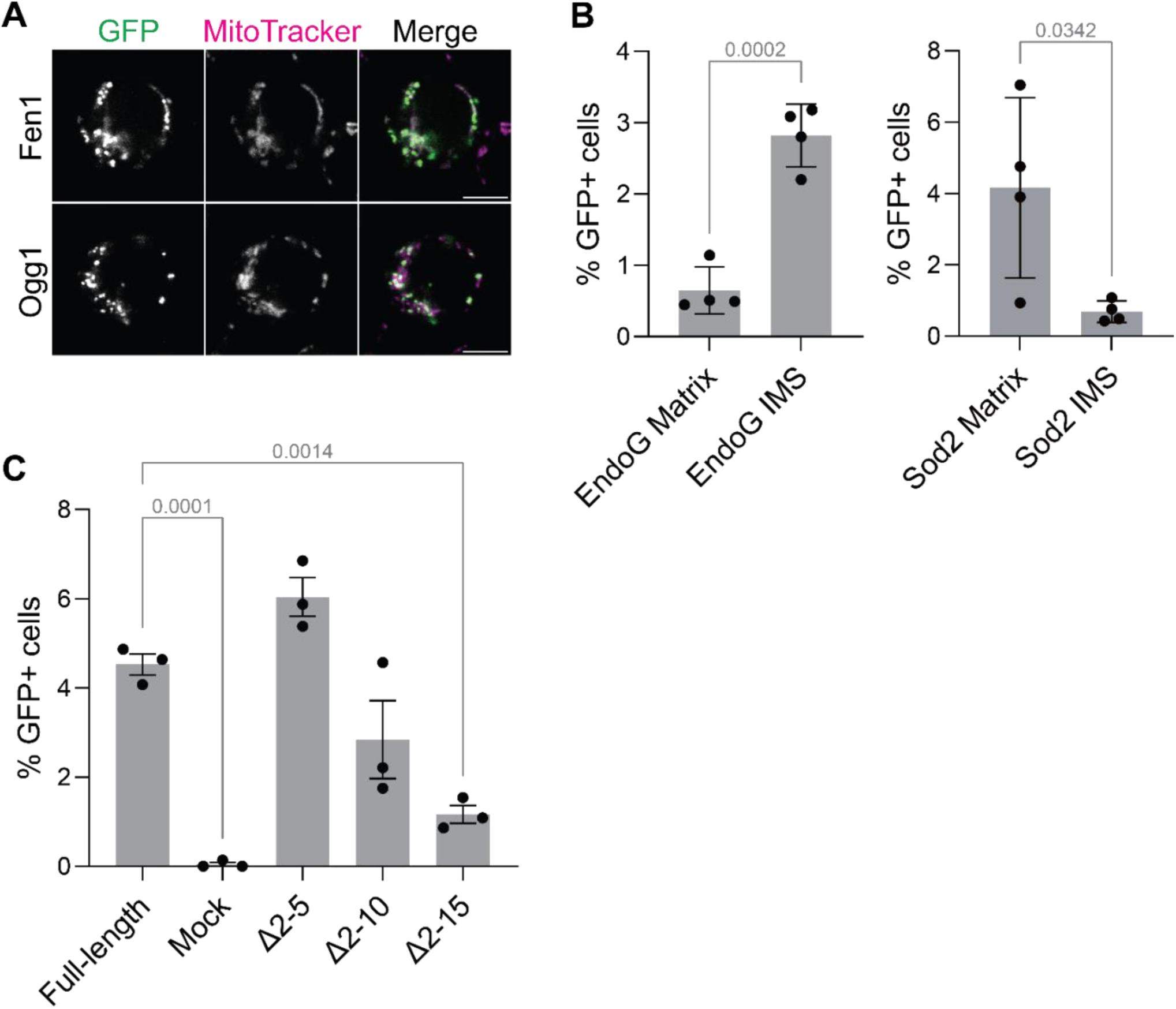
| Fen1 and Ogg1, but not EndoG, localize to the mitochondrial matrix. (**A**) Confocal micrographs of S2 cells expressing TFAM::GFP1–10 with either Fen1::GFP11×7 or Ogg1::GFP11×7. Reconstituted GFP signal colocalizes with mitochondria labelled with MitoTracker Red CMXRos. Max intensity projections of three slices across 0.595 μm. Scale bars, 5 μm. (**B**) Left: percentage of GFP-positive cells expressing TFAM::GFP1–10 (matrix) or Opa1::GFP1–10 (intermembrane space, IMS) with Sod2::GFP11×7. Right: percentage of GFP-positive cells expressing TFAM::GFP1–10 or Opa1::GFP1–10 with EndoG::GFP11×7. n = 4 independent transfections. Two-tailed Student’s t-test. Data, mean ± s.d. (**C**) N-terminal truncation analysis to define Hotaru’s mitochondrial targeting sequence (MTS). S2 cells were co-transfected with TFAM::GFP1–10 and Hotaru::GFP11×7 variants with N-terminal truncations of increasing size. n = 3 independent transfections. One-way ANOVA with Dunnett’s test. Data, mean ± s.d.

**Extended Data Fig. 2.**
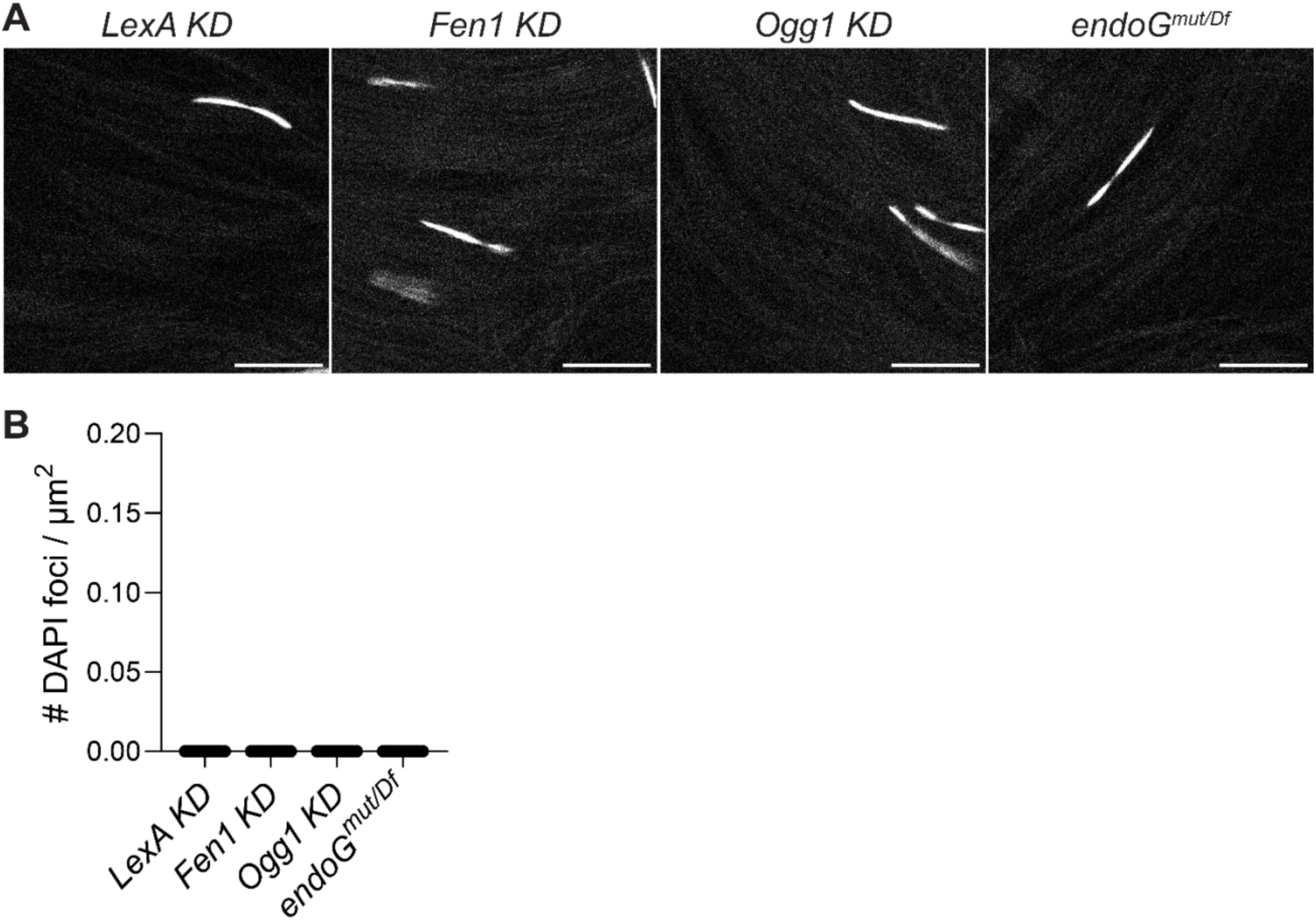
| *Fen1*, *Ogg1,* and *endoG* are not required for paternal mtDNA elimination. (**A**) Confocal micrographs of DAPI-stained sperm from males with germline knockdown of *Fen1* or *Ogg1* using *bam-GAL4* and *endoG* mutant (MB07150) heterozygous with a deficiency. (**B**) Quantification of DAPI-positive mtDNA foci for the genotypes shown in (A). n = 10 independent males. Data, mean ± s.d.

**Extended Data Fig. 3.**
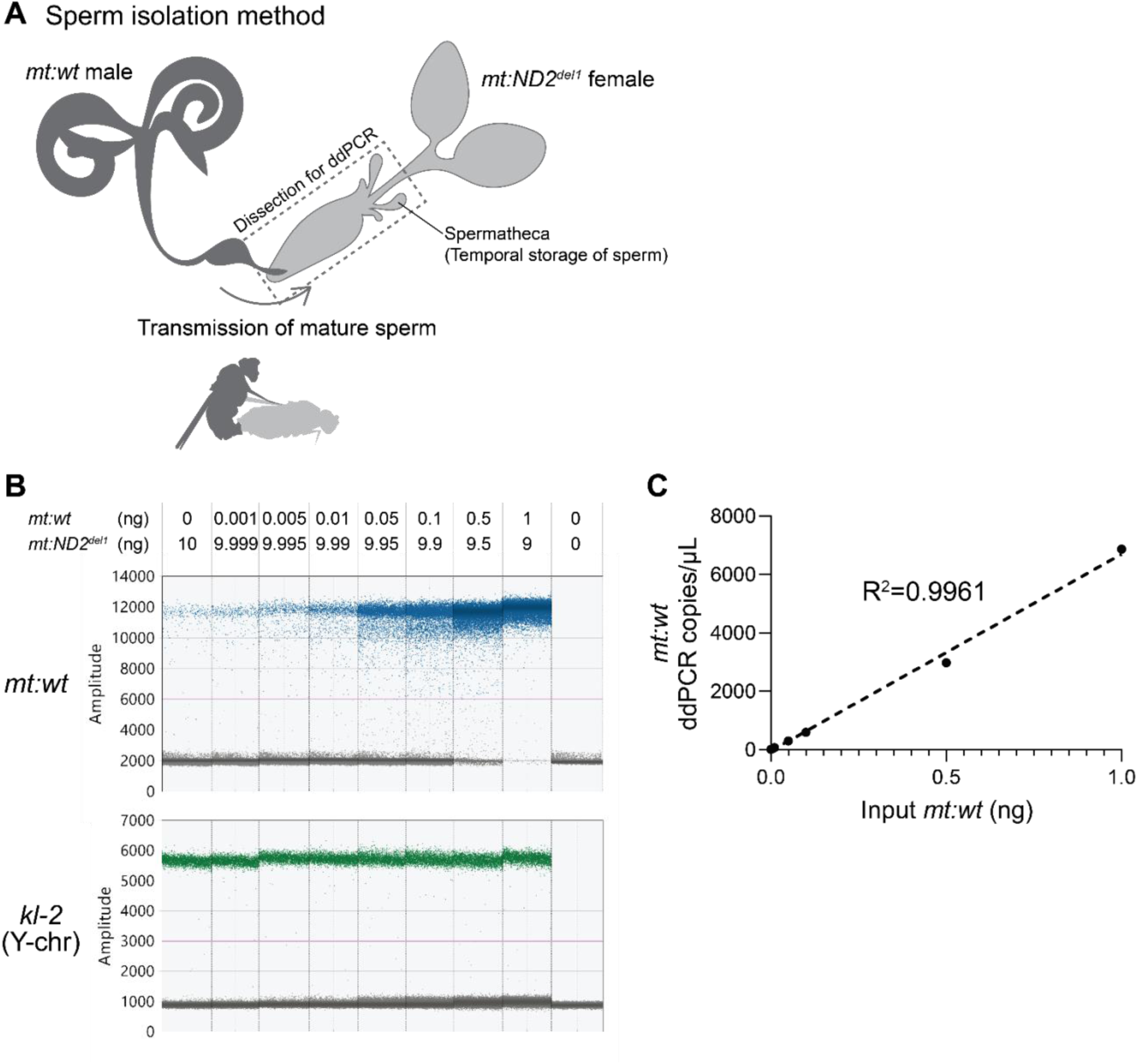
| Approach to isolate and quantify mtDNA copy number in mature sperm. (**A**) Schematic of sperm isolation strategy adapted from a previously described method^3^. Mature sperm transferred to the female during mating are stored in the spermatheca. Female reproductive tract with spermathecae was dissected, and both mtDNA copy number and sperm number were quantified by droplet digital PCR (ddPCR). To specifically detect paternal mtDNA, males carrying wild-type mtDNA (*mt:wt*) were mated to females harboring a 9-bp deletion in *mt:ND2* (*mt:ND2^del1^*). This strategy enabled the use of primers that selectively amplify paternal mtDNA while excluding maternal mtDNA. Sperm number per sample was determined by quantifying the Y chromosome (*kl-2* locus) and multiplying the measured value by two, assuming that half of sperm carry a Y chromosome. (**B**) ddPCR amplitude plot for primer/probe evaluation. Each reaction contained 0–1 ng of total DNA from *w^1118^* (*mt:wt*) flies and 9–10 ng of total DNA from *w^1118^* (*mt:ND2^del1^*) flies. *mt:wt* primers were designed to amplify *mt:wt* but not *mt:ND2^del1^*. (**C**) Linear correlation between input *mt:wt* mtDNA and mtDNA copies, quantified from (B) (R² = 0.9961).

**Extended Data Fig. 4.**
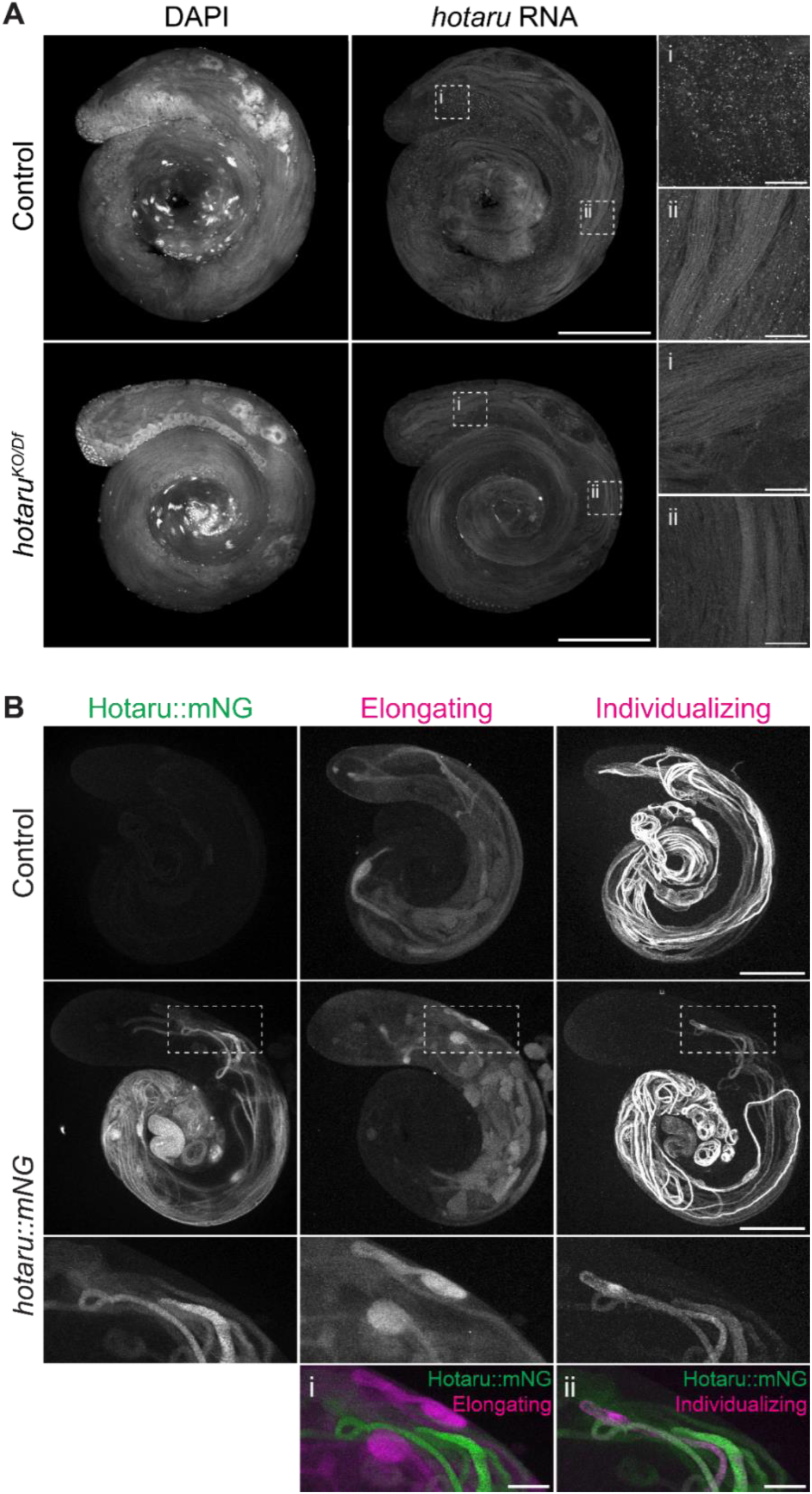
| *hotaru* RNA is expressed throughout spermatogenesis, while Hotaru protein is expressed in elongated spermatids. (**A**) HCR-FISH detection of *hotaru* RNA in control (*w^1118^*) and *hotaru^KO/Df^* testes. *hotaru* transcripts are present throughout spermatogenesis including in spermatocytes (i) and elongating/elongated spermatids (ii) in control but not in *hotaru^KO/Df^*. Scale bars: whole testis, 200 μm; insets, 20 μm. Same images for control are shown in Fig. 3C. (**B**) Confocal micrographs of control (*w^1118^*) and *hotaru::mNG* testes stained with late-elongating cyst marker anti-Soti and individualizing cyst marker anti-polyglycylated tubulin. Hotaru does not colocalize with elongating cysts (i) and colocalizes with individualizing cysts (ii). Max intensity projections across the entire thickness of testes. Scale bars: whole testis, 200 μm; insets, 50 μm. Same images for *hotaru::mNG* are shown in Fig. 3D.

**Extended Data Fig. 5.**
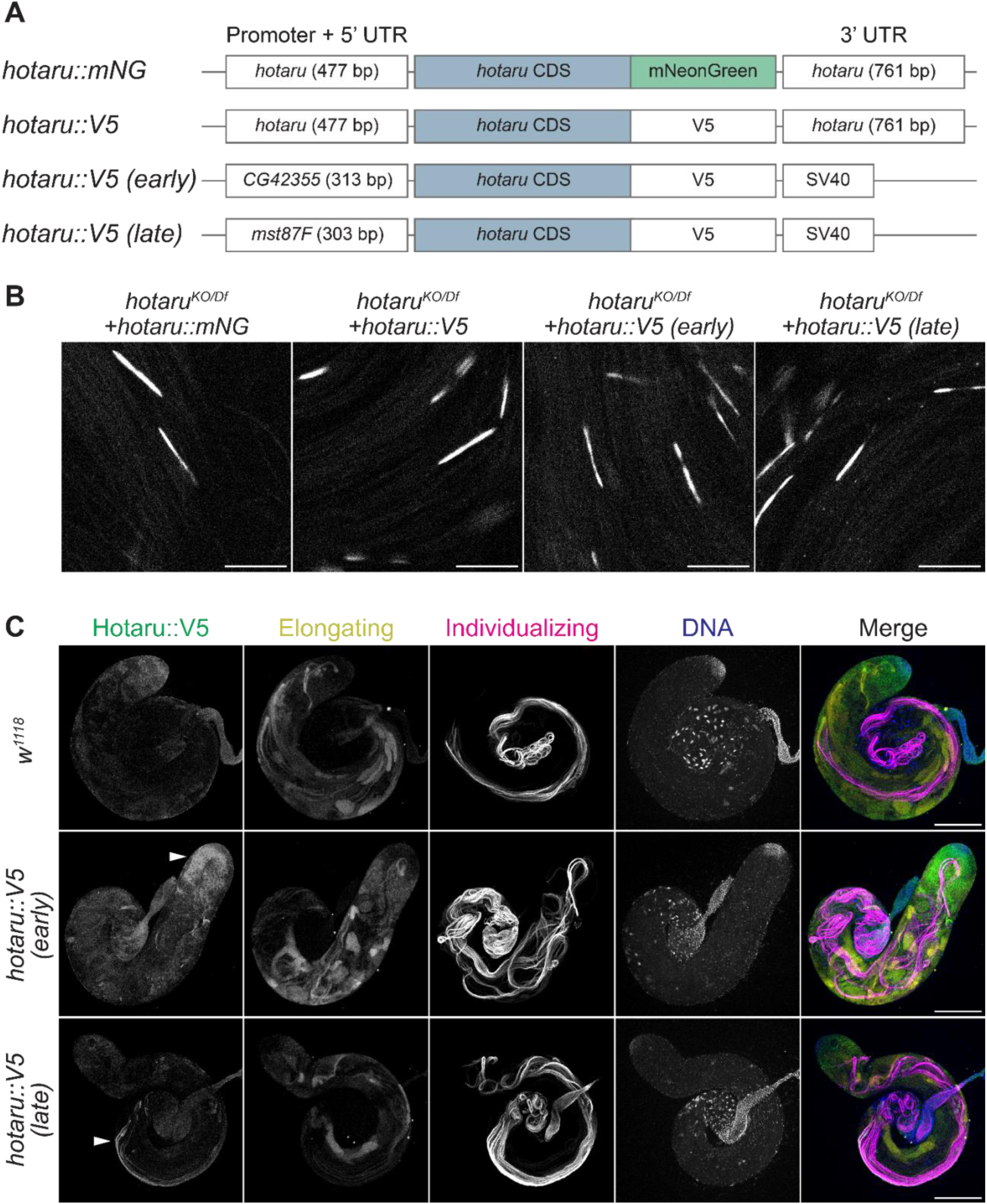
| Transgenic constructs express Hotaru at different stages of spermatogenesis. (**A**) Schematics of the transgenic constructs used in this study (not to scale). (**B**) Confocal micrographs of DAPI-stained sperm from *hotaru^KO/Df^* males expressing Hotaru under *hotaru’s* endogenous regulatory elements (positive control) or under regulatory elements of post-meiotically translated genes *CG42355* (early) and *mst87F* (late). Scale bars, 10 μm. Quantifications are shown in Fig. 3E. (**C**) Confocal micrographs of testes co-stained with anti-V5 to detect Hotaru, late-elongating cyst marker anti-Soti, and individualizing cyst marker anti-polyglycylated tubulin. Max intensity projections across the entire thickness of testes. Scale bars, 200 μm.

**Extended Data Fig. 6.**
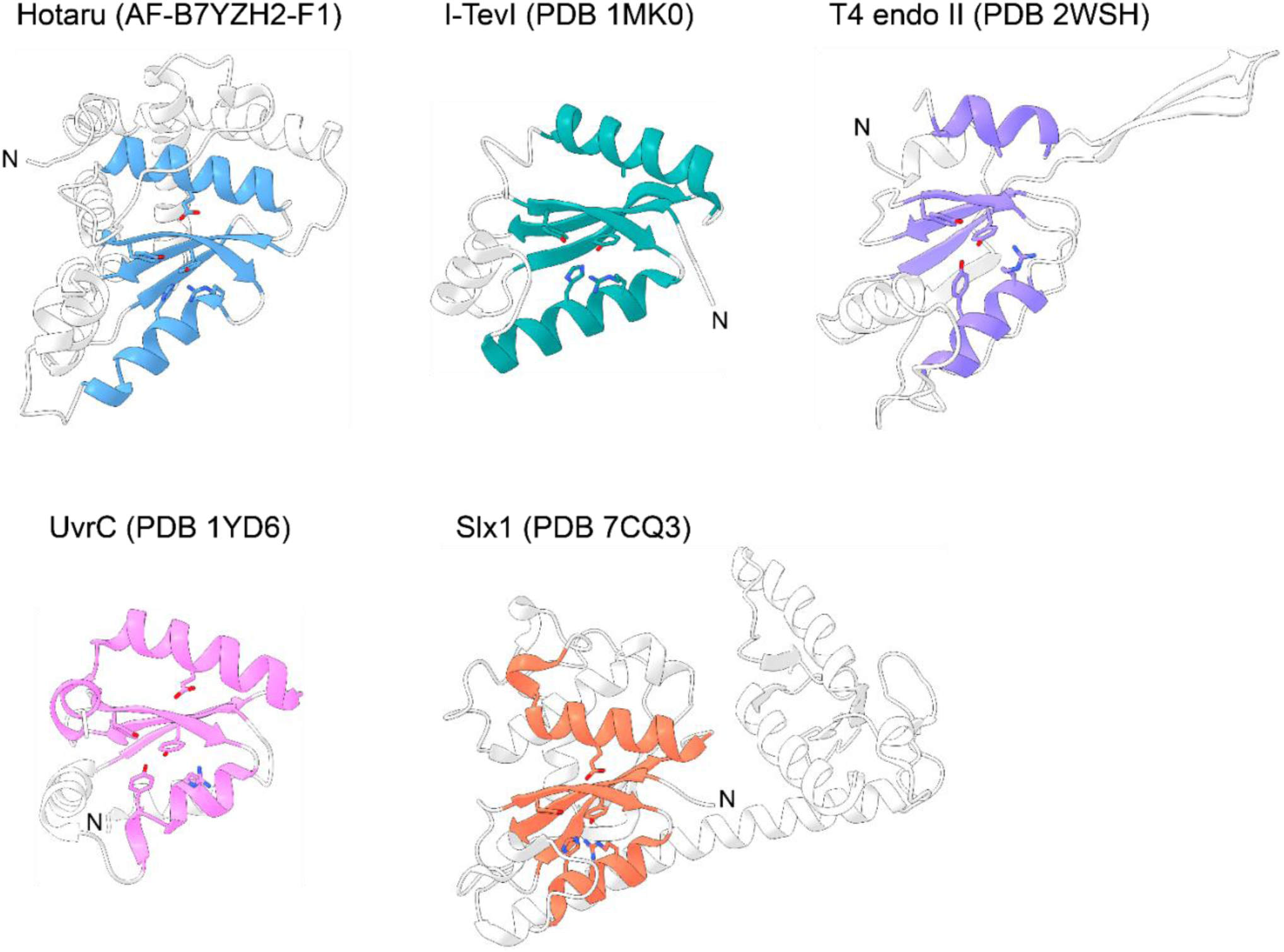
| Hotaru possesses a GIY-YIG domain. The predicted structure of Hotaru reveals a well-folded GIY-YIG domain (blue) with conserved active site residues shown as sticks. Nucleases involved in homing (I-TevI), DNA degradation (T4 endonuclease II), and DNA repair (UvrC and Slx1) contain closely related GIY-YIG catalytic cores (coloured, Table 2), characterized by a mixed α/β topology (β-β-α-β-α), in which a central three-stranded antiparallel β-sheet is flanked by α-helices^36^. The first two β-strands contain the conserved catalytic tyrosines of the GIY and YIG motifs, respectively. Additional active site residues include an invariant Arg and Glu, as well as a His that is substituted by Tyr in some lineages. In Hotaru, these correspond to R129, E186, and H133, respectively (Figure 4A). Protein Data Bank (PDB) accession codes are shown in parenthesis for experimentally determined structures; the Hotaru model is from the AlphaFold Protein Structure Database (accession code shown in parenthesis).

**Extended Data Fig. 7.**
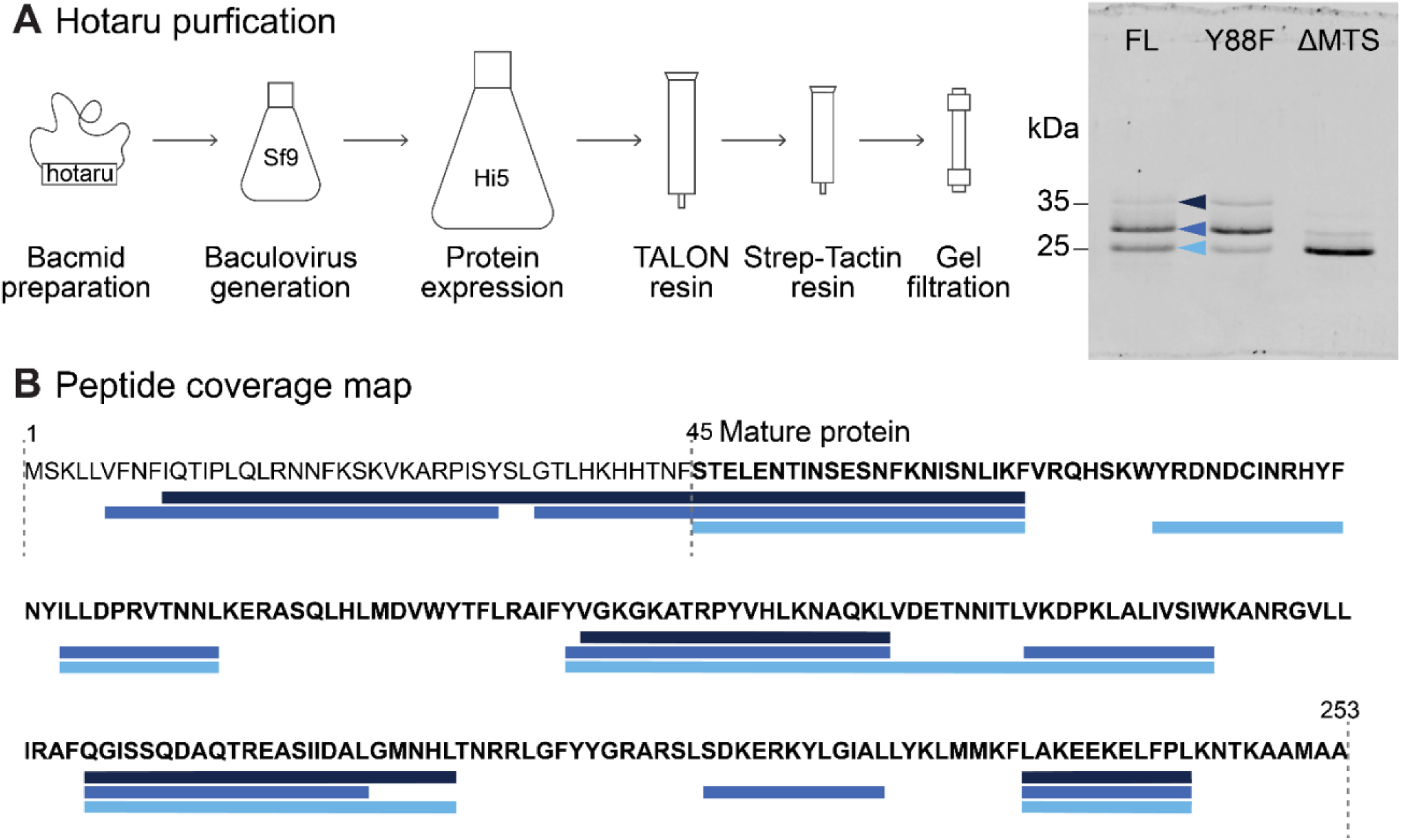
| Hotaru was purified from baculovirus-infected insect cells. (**A**) Workflow for purification of Hotaru::His10::TwinStrepII (FL, Y88F, and ΔMTS) from baculovirus-infected insect cells. Three protein species of different molecular weights were obtained from the FL and Y88F preparations. Each protein (0.5 µg) was loaded on a 12% Tris-Glycine gel and stained with InstantBlue. (**B**) Mass-spectrometry peptide coverage of the three species shown in (A), mapped onto the Hotaru protein sequence.

**Extended Data Fig. 8.**
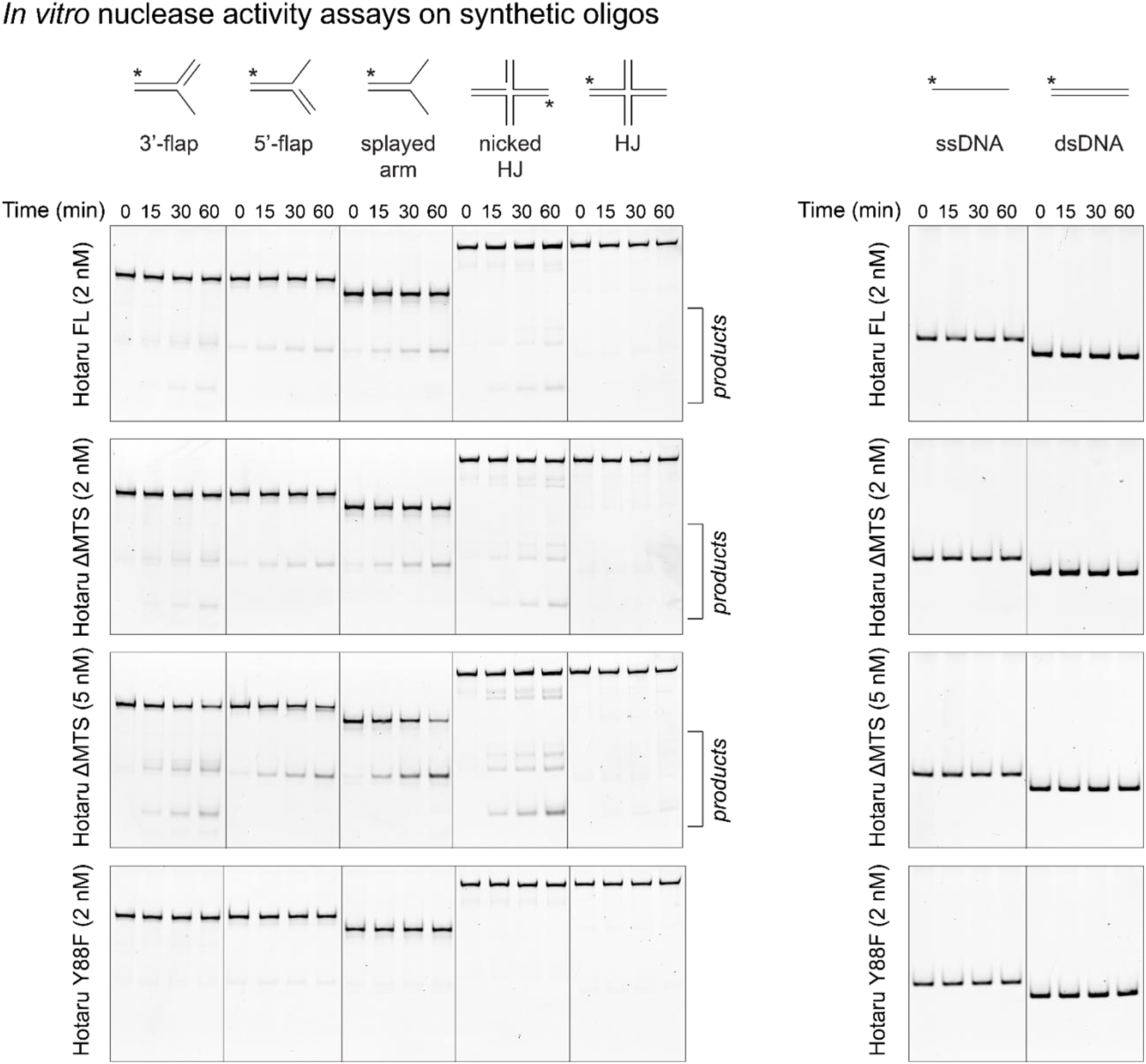
| Purified Hotaru cleaves branched DNA *in vitro*. DNA substrates (10 nM) were 5’ end-labelled with FAM on one oligonucleotide (asterisk) and incubated with purified Hotaru FL (2 nM), ΔMTS (2 nM and 5 nM), or Y88F (2 nM) at 25°C for 0, 15, 30, and 60 min. Reaction products were resolved by native PAGE.

**Extended Data Fig. 9.**
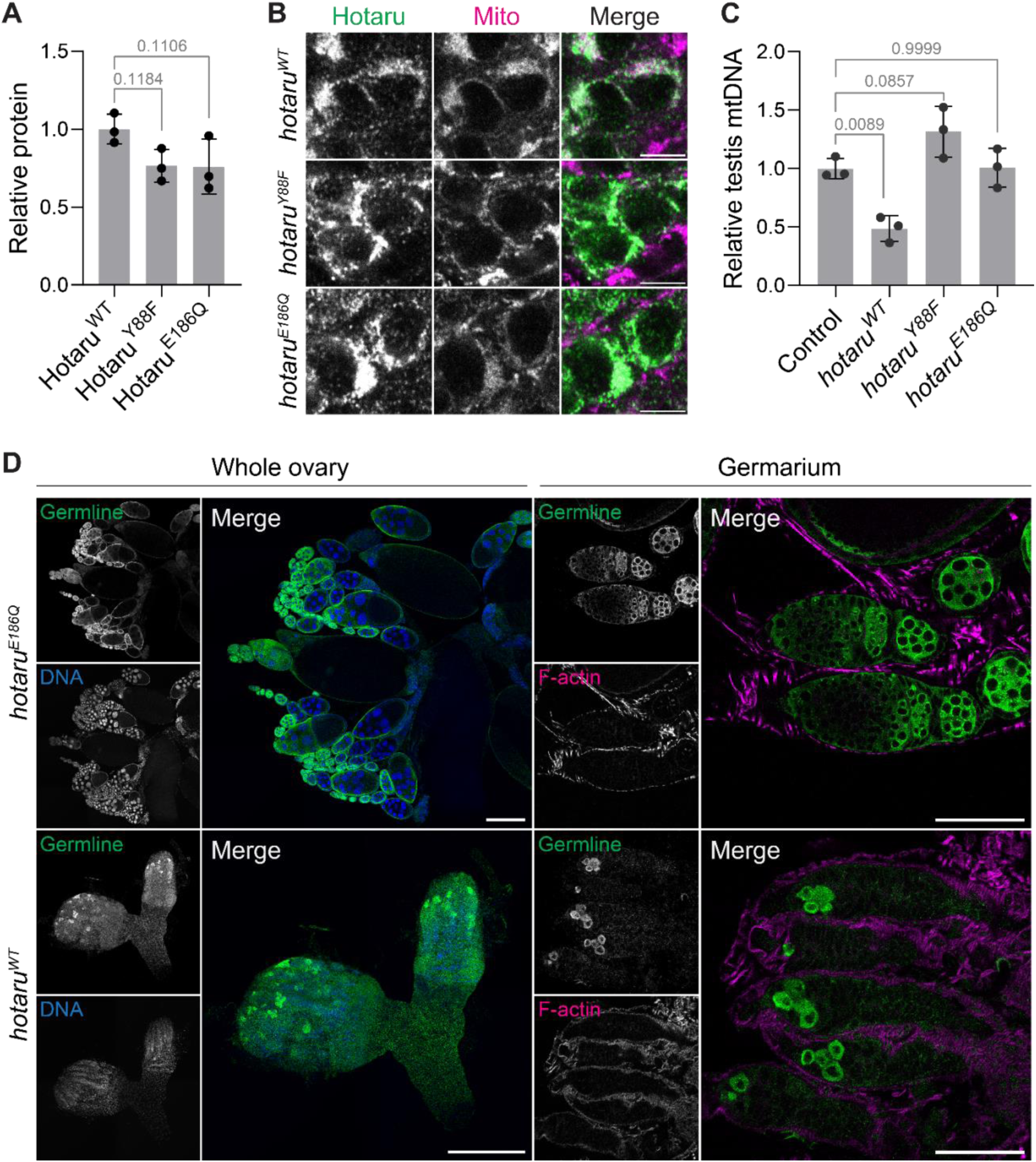
| Ectopic overexpression of Hotaru reduces mtDNA copy number. (**A**) Relative protein expression level of Hotaru::3xHA variants in S2 cells. Protein amount was quantified by Western blot and normalized to α-Tubulin. n = 3 independent transfections. One-way ANOVA with Dunnett’s test. Data, mean ± s.d. (**B**) Confocal micrographs of spermatocytes expressing *UAS-hotaru::3×HA* (*WT, Y88F,* or *E186Q*) using *bam-GAL4*. Hotaru colocalizes with mitochondria marked with anti-ATPB. Scale bars, 5 μm. (**C**) mtDNA copy number in testes expressing control (*UAS-mito::mCherry*) or *UAS-hotaru* (*WT, Y88F*, or *E186Q*) using *bam-GAL4*. mtDNA levels were measured by qPCR using mitochondrial (COX1, COX3) primers and normalized to nuclear genes (RpL11, tub). Each data point represents a pool of three pairs of testes. One-way ANOVA with Dunnett’s test. Data, mean ± s.d. (**D**) Confocal micrographs of ovaries expressing *UAS-hotaru* (WT or E186Q) in the germline stem cells using *nanos-GAL4::vp16*. Germline is marked with anti-Vasa, DNA is marked with DAPI, and F-actin is marked with phalloidin. Scale bars: whole ovary, 200 μm; germarium, 50 μm.

**Extended Data Fig. 10.**
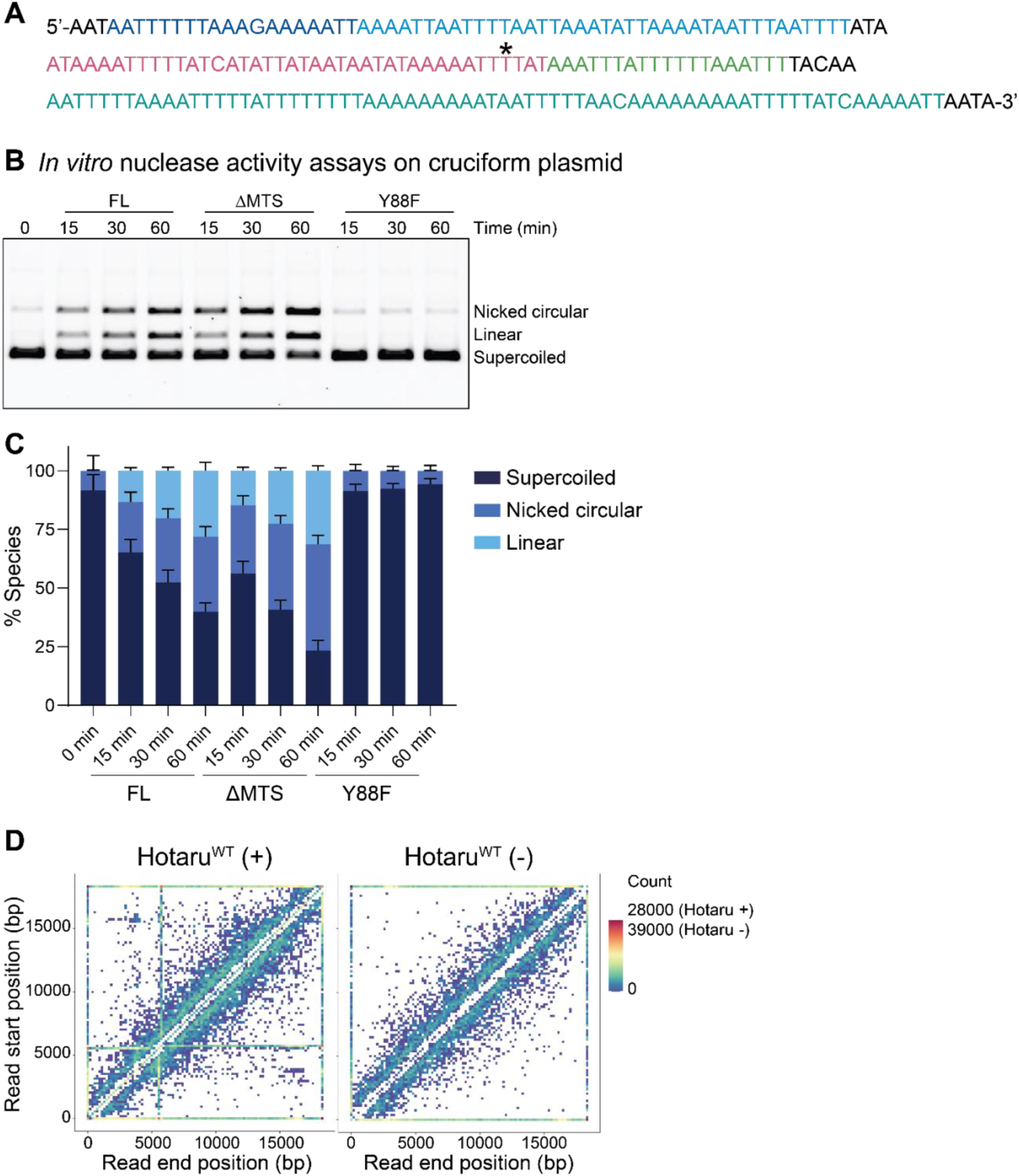
| Hotaru cleaves cruciform structures. (**A**) A prominent read start/end position (asterisk) is observed in mtDNA purified from eggs expressing *UAS-hotaru* (*WT*) using *matα-tub-GAL4*. The sequence around this position is shown and the inverted repeats are coloured. (**B**) Cruciform plasmid (pIRbke8^mut^)^45–47^ (0.5 nM) was incubated with purified wild-type Hotaru (FL or ΔMTS) or catalytically-dead Hotaru Y88F (1 nM) for 0 (no enzyme), 15, 30, and 60 min. Reaction products were resolved by 0.8% agarose gel. The cropped image for ΔMTS and Y88F is shown in Fig. 5C. (**C**) Quantification of (B). n = 4 independent assays. Data, mean ± s.d. The same quantifications are shown as line graphs for ΔMTS and Y88F in Fig. 5D. (**D**) Heat map generated from ONT reads of cruciform plasmid with or without 60 min incubation with Hotaru. Hotaru-treated plasmid has a cleavage hotspot in the inverted repeat. Same heat map for Hotaru-treated sample is shown in Fig. 5E.

**Extended Data Fig. 11.**
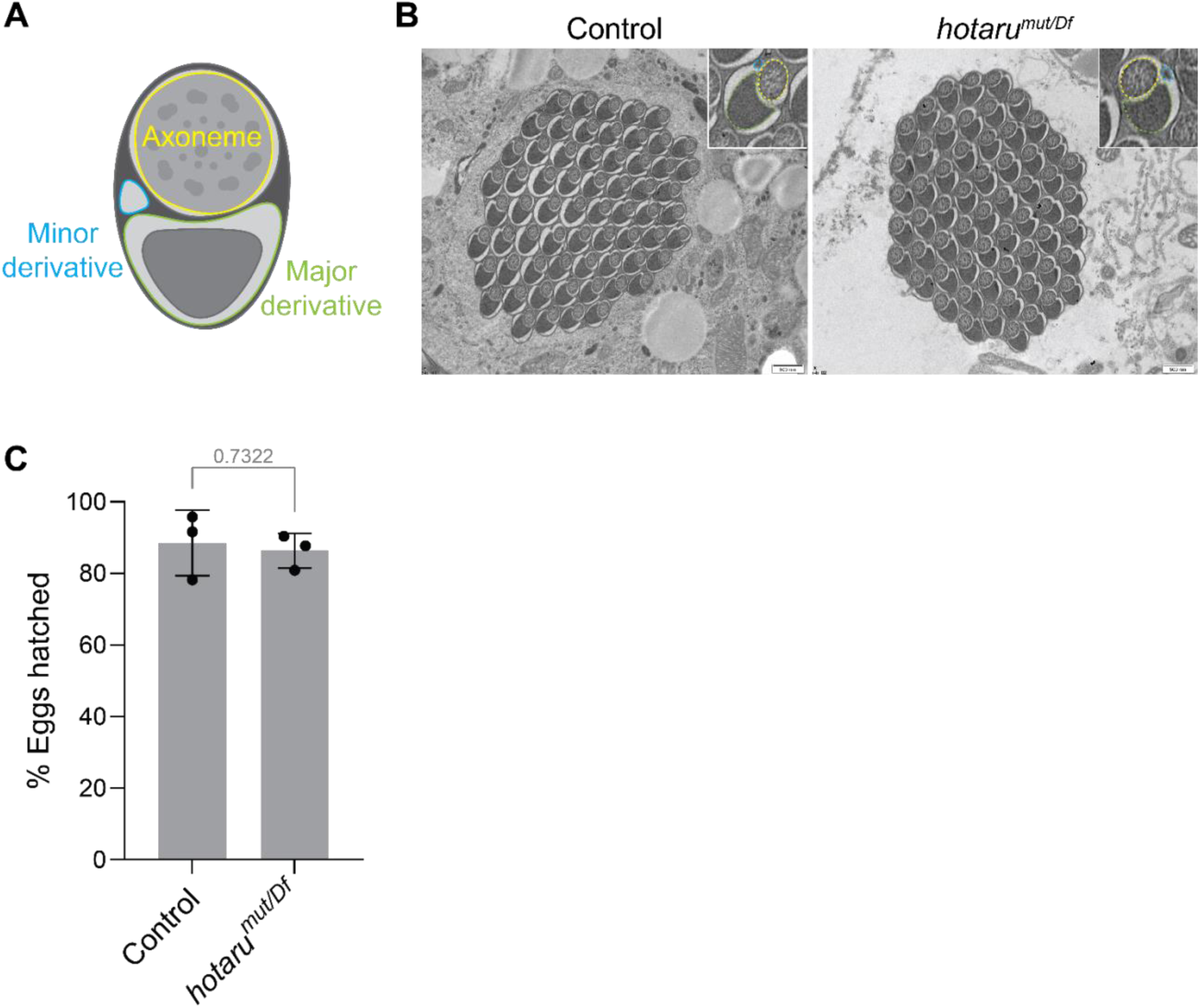
| *hotaru* mutant sperm develop normally and are fertile. (**A**) Cartoon illustrating a vertical section of an individualized spermatid. Two mitochondrial derivatives extend along the axoneme throughout the entire length of the spermatid. (**B**) Transmission electron micrographs of individualized spermatid cysts from control (*w^1118^*) and *hotaru^mut/Df^* males. *hotaru^mut/Df^* spermatids are morphologically normal. Scale bars, 500 nm. (**C**) Percentage of eggs hatched when fertilized by control (*w^1118^*) or *hotaru^mut/Df^* males. Each data point represents eggs laid by 5 females mated with 3 males, averaged over 3 days. Two-tailed Student’s t-test. Data, mean ± s.d.

## METHODS

### Fly husbandry

All flies were reared on standard medium (cornmeal, yeast, agar, and molasses) at 25°C and 60%–70% humidity on a 12-hour light/12-hour dark cycle.

### Split-GFP experiments

For the split-GFP experiments, the coding sequence of endonucleases and TFAM were PCR amplified from wild-type (*w^1118^*) cDNA. We used FlyBase^51,52^ (release FB2023_06) to obtain the sequences. TFAM was fused C-terminally to GFP1–10 and a 3×HA tag, whereas the endonucleases were fused C-terminally to seven tandem GFP11 repeats and a 3×FLAG tag. Constructs were cloned into the pAc5.1 expression vector using NEBuilder HiFi DNA Assembly Master Mix (NEB, E2621L).

S2 cells were cultured in Express Five SFM (Gibco, 10486025) supplemented with 20 mM L-glutamine (Sigma-Aldrich, G7513) and Penicillin-Streptomycin (Sigma-Aldrich, P4333). S2 cells were transfected with GFP1-10 and GFP11x7 plasmids using Effectene transfection reagent (Qiagen, 301425). Two days later, 200 µL cells were transferred to a well of a live imaging chamber (ibidi, 80806) and imaged using a Leica SP8 confocal microscope. To quantify the % GFP-positive cells, three fields per well were imaged with a 20x, NA 0.75 dry objective lens and the total number of cells and the number of GFP-positive cells were automatically counted using the macros function on ImageJ. Values from the three fields were averaged and represented as one data point. For imaging individual GFP-positive cells, mitochondria were labelled with MitoTracker Red CMXRos (25 nM) and imaged with a 63x, NA 1.4 immersion oil objective lens.

### S2 cell immunofluorescence

For immunofluorescence, S2 cells were transfected with various truncated versions of Hotaru C-terminally tagged with 3xFLAG using Effectene transfection reagent (Qiagen, 301425), directly in the imaging chamber (ibidi, 80806). Two days later, cells were fixed with 4% formaldehyde for 15 min, permeabilized and blocked in PBSTB (PBS, 0.1% Triton X-100, 1% BSA) for 30 min, and incubated with primary antibody (anti-FLAG 1:1000 and anti-ATP5A 1:1000) in PBSTB for overnight at 4°C. The next day, cells were incubated with secondary antibody (anti-Mouse IgG1-Alexa 488 1:500 and anti-Mouse IgG2b-Alexa 568 1:500) for 30 min and stained with 1 µg/mL DAPI in PBS. Images were taken with a 63x, NA 1.4 immersion oil objective lens.

### S2 cell immunoblotting

To measure the protein levels of Hotaru variants in S2 cells, Hotaru::3xHA (WT, Y88F, or E186Q) was cloned into the pAc5.1 vector, and transfected into S2 using Effectene transfection reagent (Qiagen, 301425). Two days later, S2 cells were pelleted, frozen, directly resuspended in 5x Laemmli buffer supplemented with 5% 2-mercaptoethanol, and boiled before loading onto 4–15% Precast Protein Gel (Bio-Rad, 4561086). The blot was detected with primary antibody (anti-HA 1:5000 or anti-α-Tubulin 1:5000) followed by secondary antibody (anti-Rabbit-HRP 1:10000 or anti-Mouse-HRP 1:10000). The blots were imaged using an iBright 1500 imager (Invitrogen) and quantified using ImageJ.

### Transgenic and CRISPR flies

For the deletion of the *hotaru* locus, the following oligos were synthesized (IDT) and cloned in tandem into the pCFD5_w vector (Addgene, #112645): TAAGCTGCAAAGGGATGGTT (gRNA 1), AAGTTATTTCTAAGCTGCAA (gRNA 2), TGGTCGAGCTCGAAGTTTAT (gRNA 3), and AGCTCGAAGTTTATCGGATA (gRNA 4). Homology arms containing ∼1 kb upstream and ∼0.9 kb downstream of *hotaru* was cloned into pScarlessHD-DsRed-w+ (Addgene, #80801). gRNA and homology donor plasmids were injected into Vasa-Cas9 (BL51323) expressing embryos (Genome Prolab). Eclosed flies were crossed to *w^1118^* flies, and the progeny with successful knockout were screened by the presence of DsRed. For *UAS-hotaru*, Hotaru was C-terminally tagged with 3xHA and cloned into pUASz^53^. For genomic *hotaru* transgenes, the region spanning from 477 bp upstream to 761 bp downstream of the *hotaru* CDS was cloned into pCaSpeR4, and mNeonGreen or V5 tag was inserted at the 3’ end of the *hotaru* CDS. For post-meiotic expression constructs, from the genomic *hotaru::V5* transgene, *hotaru*’s 5’UTR was changed to the 5’UTR of *CG42355* (313 bp upstream of CDS) or *mst87F* (303 bp upstream of CDS), and *hotaru*’s 3’UTR was changed to SV40. Sanger sequencing (The Centre for Applied Genomics) or whole amplicon sequencing (Plasmidsaurus) was performed to verify the genotypes of all transgenic and CRISPR flies.

### DAPI staining of mature sperm

0 to 1-day-old males were separated from females for three days to accumulate sperm in their seminal vesicles. Seminal vesicles were dissected, transferred to a PBS droplet on a poly-L-lysine-coated slide (Electron Microscopy Sciences, 63410), and sandwiched under a coverslip. PBS was removed from the side of the coverslip with Kimwipes to flatten the seminal vesicle and extrude the sperm. The slide was snap frozen in liquid nitrogen and the coverslip was popped off with a razor blade. Fluoromount-G mounting medium with DAPI (Invitrogen, 00-4959-52) was added to the sample area, sealed with a coverslip, and fixed with nail polish. Samples were imaged with a Leica SP8 confocal microscope using a 63x, NA 1.4 immersion oil objective lens.

### Relative quantification of DAPI-stained mtDNA in sperm

The number of DAPI-positive puncta was quantified using the Imaris (Oxford Instruments, v9.8) Spots tool and normalized to the sperm occupying area measured by ImageJ. The quantified values should be interpreted solely as relative estimates of mtDNA abundance and not absolute mtDNA copy number per sperm.

### ddPCR assay design and evaluation

Primers and probes were designed based on previous work^3^ with some modifications. To specifically detect paternal wild-type mtDNA (*mt:wt*) without detecting maternal mtDNA carrying the 9-bp deletion (*mt:ND2^del1^*), a primer pair and a FAM-labelled probe were designed at this region. Additionally, another primer pair and a HEX-labelled probe targeting a Y-chromosome gene *kl-2* were used to quantify the sperm numbers. Primers/probe sets (IDT) used in the assay are listed in Table 1. The assay was performed on the QX200 Droplet Digital PCR system (Bio-Rad), with each 20 µL reaction containing 10 µL of 2× ddPCR Supermix for Probes (No dUTP) (Bio-Rad, 1863024), 1 µL of the mt:ND2 assay (FAM-labelled), 1 µL of the kl-2 assay (HEX-labelled), and DNA. The assay was validated using a temperature gradient to ensure optimal separation of target and reference droplets. Thermal cycling was performed on a Veriti thermal cycler (Thermo Fisher Scientific) with the following conditions: 95 °C for 10 min, followed by 45 cycles of 94 °C for 30 s and 50 °C for 1 min, then 98 °C for 10 min, and a final hold at 10 °C. All steps used a ramp rate of 2 °C/s. Data were analyzed using QuantaSoft v1.7.4 (Bio-Rad).

**Table 1.**
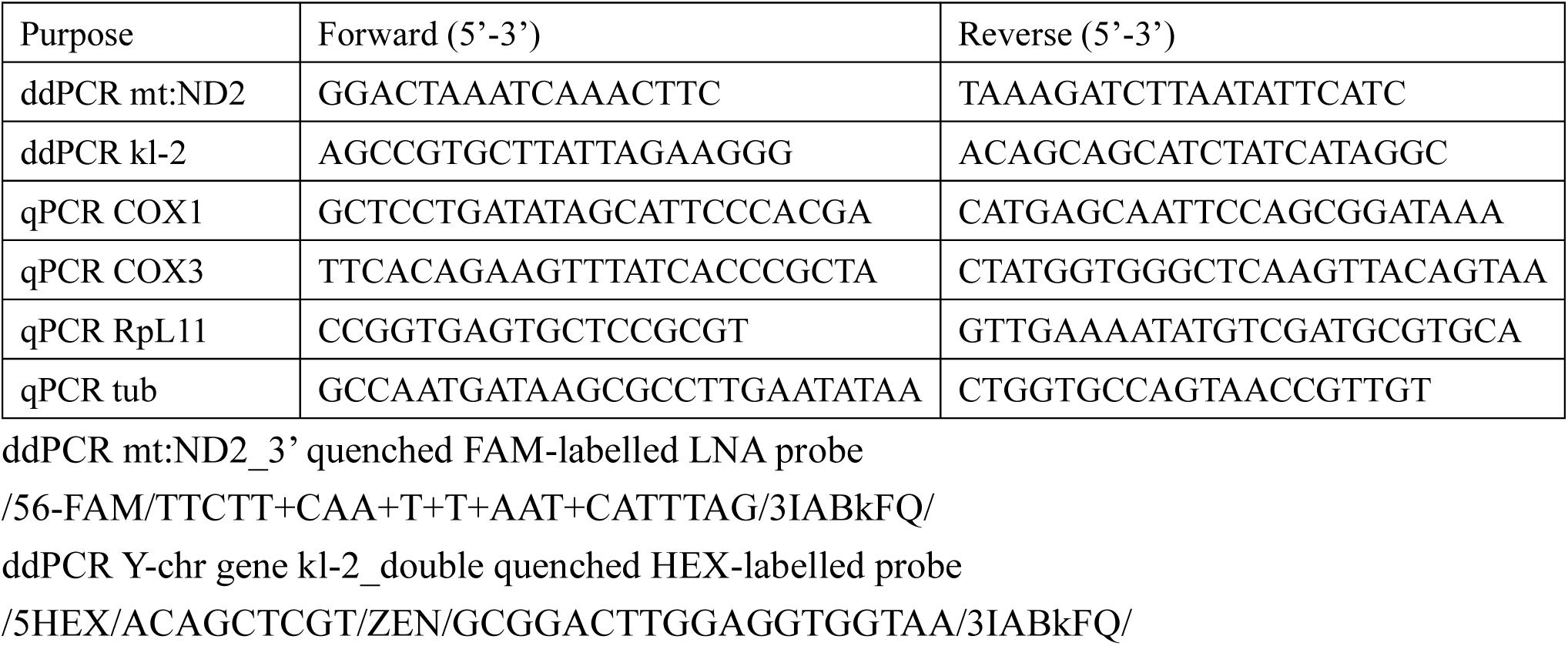
qPCR/ddPCR primers/probes used in this study.

**Table 2.** DALI structural similarity search results for the Hotaru LEM-3-like GIY-YIG domain (residues 85-244).

| PDB and Chain | Z-Score | RMSD (Å) | Number of Aligned Residues (LALI) | Number of Residues in the Structure (NRES) | Sequence Identity (%) |
| --- | --- | --- | --- | --- | --- |
| 1MK0-A | 7.8 | 2.0 | 84 | 97 | 15 |
| 2WSH-A | 5.8 | 2.7 | 83 | 134 | 17 |
| 1YD6-D | 4.8 | 2.2 | 69 | 96 | 20 |
| 2LWF-A | 3.4 | 2.4 | 64 | 119 | 14 |

To evaluate the specificity of the primers/probe set for *mt:ND2*, varying amounts (0, 0.001, 0.005, 0.01, 0.05, 0.1, 0.5, 1 ng) of genomic DNA of *w^1118^* (*mt:wt*) flies were mixed with genomic DNA of *w^1118^* (*mt:ND2^del1^*) flies to total amounts of 10 ng. Genomic DNA was isolated using the DNeasy Blood and Tissue Kit (Qiagen, 69504). The coefficient of correlation (R^2^) was calculated to be 0.9961, showing reliable correlation between the input *w^1118^* (*mt:wt*) DNA amount and the measured copy number.

### mtDNA quantification in mature sperm

Experiment was performed similarly to previously described^3^. 20 2-day-old *mt: ND2^del1^* virgin females were crossed to 15 2-day-old males of experimental genotypes for 24 h at 25 °C (Extended Data Fig. 3A). The mated females were rinsed with 75% ethanol on the gas pad to remove any male body parts. The reproductive tract, including the spermathecae and the seminal receptacle, but excluding the ovary, was dissected in ice-cold PBS and transferred into microfuge tubes on ice using a glass pipette. Samples were spun down and PBS was removed before storing them at -80°C. To process samples for ddPCR, 100 µL of high-Tris squishing buffer (100 mM Tris-HCl pH 8.0, 100 mM NaCl, 25 mM EDTA) with 2% SDS and 200 µg/mL proteinase K (NEB, P8107S) was added to the pellet and incubated at 56°C for 2 h with occasional vortexing. Then, DTT was added to 0.2 M and incubated at 99°C for 10 min. Samples were brought back to room temperature and subjected to DNA extraction by phenol-chloroform followed by ethanol precipitation. The DNA pellet was resuspended in 20 µL double-distilled water, and 4 µL was used as input for ddPCR. mtDNA copy number per sperm was calculated by dividing mt:*ND2* (mtDNA) copies by twice the number of *kl-2* (Y-chromosome) copies. All genotypes were created in the same mtDNA background to avoid mismatch between DNA template and primers/probes.

### mtDNA quantification in testes and ovaries

Three pairs of testes or ovaries were collected in 50 µL (testis) or 100 µL (ovary) Squishing buffer (10 mM Tris-HCl pH 8.2, 25 mM NaCl, 1 mM EDTA) with 0.2% SDS and 200 µg/mL proteinase K (Bioshop, PRK403.100). Tissues were homogenized using plastic pestles, incubated at 55 °C for 25 min, and heat-inactivated at 97 °C for 5 min. For qPCR, 0.2 µL of the lysate was used in a 10 µL reaction containing SensiFAST SYBR No-ROX (Bioline, BIO-98050). mtDNA levels (COX1, COX3 primers) were normalized to nuclear DNA levels (RpL11, tub primers).

### Immunostaining of Drosophila testes

Immunostaining was performed as previously described^31^. Testes were dissected from 0 to 1-day-old males in ice-cold TB (10 mM Tris-HCl pH 6.8, 183 mM KCl, 47 mM NaCl, 1 mM EDTA) and transferred to a drop of TB on a poly-L-lysine-coated slide (Electron Microscopy Sciences, 63410). Testes were sandwiched flat between the slide and a coverslip, excess TB was removed with Kimwipe, and snap frozen in liquid nitrogen. The coverslip was popped off with a razor blade, and the samples were plunged into -20°C ethanol, for at least 10 min. Samples were fixed with 4% formaldehyde in PBS for 7 min, rinsed with PBT (PBS, 0.1% Triton X-100), permeabilized in PBT DOC (PBS, 0.3% Triton X-100, 0.3% Sodium deoxycholate) for 15 min twice, washed with PBT, and blocked in PBTB (PBS, 0.1% Triton X-100, 5% BSA) at RT for 30 min. Samples were stained with primary antibodies (anti-mNeonGreen 1:500 or anti-V5 1:500, anti-Polyglycylated tubulin 1:2500, and anti-Soti 1:100) in PBTB at 4°C overnight, washed with PBTB, and stained with secondary antibodies (1:500) in PBTB at RT for 1 h. Then, they were washed with PBT, stained with 1 µg/mL DAPI in PBT for 15 min, washed with PBT again, and mounted in Vectashield Plus mounting medium (Vector Labs, H-1900-2).

### Immunostaining of Drosophila ovaries

Females aged 0–1 days were yeasted overnight, and ovaries were dissected in PBS. Ovaries were fixed in 4% formaldehyde in PBS for 15 min, permeabilized in PBST (PBS, 1% Triton X-100) for 30 min, and blocked in PBSTB (PBS, 0.1% Triton X-100, 1% BSA) for 1 h. Samples were incubated overnight at 4°C with anti-Vasa antibody (1:4000) in PBSTB, washed in PBSTB, and incubated with anti-Rabbit-Alexa 488 (1:500) and Phalloidin-Alexa 568 (1:250) in PBSTB for 2 h at room temperature. Ovaries were then washed in PBST, stained with 1 µg/mL DAPI in PBST for 10 min, washed again in PBST, and mounted in Vectashield Plus mounting medium (Vector Labs, H-1900-2).

### HCR-FISH

HCR-FISH was performed similarly to previously described^54^. Testes were dissected from 0 to 1-day-old males in PBT (PBS, 0.1% Tween-20) and fixed in 4% formaldehyde in PBT for 20 min. Samples were washed in PBT and permeabilized overnight in 70% ethanol at -20°C. Hybridization and hairpin amplification were performed according to the manufacturer’s protocol (Molecular Instruments). Briefly, testes were washed in 5× SSCT and incubated in 50 µL probe solution (4 nM probe in hybridization buffer) at 37 °C overnight (≥16 h). After washes in wash buffer and 5× SSCT, samples were incubated in 50 µL hairpin solution (58 nM each of hairpins h1 and h2) overnight at room temperature in the dark (≥16 h). Testes were washed in 5× SSCT, stained with 1 µg/mL DAPI in SSCT for 10 min, washed in PBS, and mounted in Vectashield Plus mounting medium (Vector Labs, H-1900-2).

### Transmission electron microscopy (TEM)

Testes from 0 to 3-day-old adult flies were dissected in ice-cold PBS and immediately transferred into ice-cold Trump’s fixative (Electron Microscopy Sciences, 11750), where they were kept for 2h. Samples were transferred back to PBS, post-fixed with 1% OsO4 for 1 h, rinsed and dehydrated with an acetone series and embedded in Epon resin. Images were acquired with a Hitachi HT7800 TEM and an EMSIS Xarosa camera (The Hospital for Sick Children, Nanoscale Biomedical Imaging Facility).

### Male fertility assay

On day 0, three 2-day-old males of the specified genotypes were paired with five virgin *w^1118^* females in a cage with a apple juice agar plate with yeast paste. On days 3–5, the apple plate was exchanged every 24 h and the number of eggs laid was counted. Hatched eggs were counted 24–48 h after egg lays. The hatching rate, which is the proportion of hatched eggs among the total eggs laid, was averaged over the three days and represented as one data point.

### Purification of Hotaru

#### Cloning

Hotaru::His10::TwinStrepII construct was generated by cloning the *hotaru* coding sequence (DmeI_CG42391; Uniprot ID B7YZH2) fused with His10::TwinStrepII tag optimized for expression in *Spodoptera frugiperda* (Sf9) (Twist Bioscience) into the pFL vector^55^. Hotaru ΔMTS::His10::TwinStrepII and Hotaru Y88F::His10::TwinStrepII were generated from Hotaru::His10::TwinStrepII by amplifying the region excluding the coding sequence for amino acids 2-41 or introducing a point mutation, respectively, and performing a single-piece Gibson reaction (NEB, E2621L).

#### Baculovirus generation

Bacmid DNA was generated for Hotaru::His10::TwinStrepII (FL, ΔMTS, and Y88F) by transforming MultiBac *Escherichia coli* with the respective plasmid DNA^55^. Transformants were selected on LB plates containing kanamycin (50 μg/mL), gentamycin (7 μg/mL), and tetracycline (10 μg/mL), bluo-gal (100 μg/mL), and IPTG (40 μg/mL). Positive clones were identified using blue-white screening. Bacmid DNA was purified using the NucleoBond BAC 100 Kit (Macherey-Nagel, 740579), according to the manufacturer’s instructions, and stored at 4°C.

To produce passage 1 (P1) baculovirus, 1,000,000 Sf9 cells were transfected with 1 μg of bacmid DNA using FuGENE HD Transfection Reagent (Promega, E2311). Briefly, 3 μL of transfection reagent was diluted in 100 μL iMax serum-free media (Wisent Inc., 301-045), to which 1 μg of bacmid DNA was added. The transfection mixture was incubated at room temperature for 20 min and then added dropwise to the Sf9 cells, and incubated for 24 h at 27°C in ambient CO_2_, before replacing the growth media with fresh iMax serum-free media.

P1 baculovirus was harvested 72 h post-transfection by centrifuging the growth media at 1,800 g for 5 min (4°C), passing through a 0.2 μm Acrodisc syringe filter (MilliporeSigma), and supplementing with fetal bovine serum (10% final concentration) (Wisent Inc.) and HyClone Amphotericin B (2.5 μg/mL final concentration) (Cytiva). Viral titre was determined using the baculoQuant One-Step Titration Kit (Genway Biotech), as per the manufacturer’s instructions.

P2 baculovirus was generated by infecting Sf9 cells with P1 virus at a multiplicity of infection (MOI) of ≤ 0.1. The infected cell culture was counted every 24 h and maintained at a cell density of 1,000,000 cells/mL. Cells were harvested 48 h after proliferation had ceased, and baculovirus was collected and titrated as described above. All baculovirus stocks were stored at 4°C and protected from light.

#### Protein purification

Hotaru::His10::TwinStrepII (FL, ΔMTS, and Y88F) was purified from *Trichoplusia ni* (Hi5) cells infected with P2 baculovirus for 72 h at 27°C in ambient CO_2_. Cells were pelleted by centrifugation and washed once with cold PBS. Approximately 4 g of cells were resuspended in 40 mL lysis buffer (50 mM HEPES pH 7.6, 500 mM NaCl, 10% glycerol, 0.5% sarkosyl, 5 mM imidazole) supplemented with protease inhibitors (10 μg/mL aprotinin, 4 μg/mL bestatin, 10 μg/mL leupeptin, 10 μg/mL pepstatin, 1 mM PMSF, 1 mM benzamide) and phosphatase inhibitors (10 mM β-glycerophosphate, 5 mM sodium fluoride, 1 mM sodium orthovanadate, 1 mM sodium pyrophosphate). We found that sarkosyl was uniquely required for the efficient solubilization of Hotaru::His10::TwinStrepII. The cell suspension was homogenized on ice with pestle A (10 strokes) and then pestle B (10 strokes). Micrococcal nuclease (20 U/mL) (NEB) and CaCl_2_ (5 mM) were added to digest chromatin and nucleic acids and reduce viscosity. The lysate was incubated on ice for 30 min and then clarified by centrifugation for 1 h at 12,000 g at 4°C.

The soluble extract was incubated with 1 mL pre-equilibrated TALON Superflow resin (Cytiva, 28957499) for 2 h on a spinning wheel at 4°C. The sample was subsequently transferred to an Econo-Pac chromatography column (Bio-Rad). After washing the resin once with 5 CV lysis buffer, the resin was washed twice with 5 CV TALON wash buffer 1 (50 mM HEPES pH 7.6, 500 mM NaCl, 10% glycerol, 0.5% sarkosyl, 10 mM imidazole), TALON wash buffer 2 (50 mM HEPES pH 7.6, 500 mM NaCl, 10% glycerol, 0.1% sarkosyl, 10 mM imidazole), and TALON wash buffer 3 (50 mM HEPES pH 7.6, 500 mM NaCl, 10% glycerol, 0.1% sarkosyl, 25 mM imidazole). Each wash was incubated for 5 min at 4°C. Bound proteins were eluted sequentially using 2 x 2 CV TALON elution buffer 1 (50 mM HEPES pH 7.6, 500 mM NaCl, 10% glycerol, 0.1% sarkosyl, 50 mM imidazole), TALON elution buffer 2 (50 mM HEPES pH 7.6, 500 mM NaCl, 10% glycerol, 0.1% sarkosyl, 100 mM imidazole), and TALON elution buffer 3 (50 mM HEPES pH 7.6, 500 mM NaCl, 10% glycerol, 0.1% sarkosyl, 250 mM imidazole). Each elution was incubated for 5 min at 4°C. Peak fractions were identified by SDS-PAGE, pooled, and then incubated with 0.4 mL pre-equilibrated Strep-TactinXT 4Flow high-capacity resin (IBA Lifesciences, 2-5030-010) for 10 min at 4°C. The resin was washed twice with 5 CV Strep wash buffer 1 (50 mM HEPES pH 7.6, 500 mM NaCl, 10% glycerol, 0.1% sarkosyl, 1 mM DTT) and then twice with 5 CV Strep wash buffer 2 (50 mM HEPES pH 7.6, 250 mM NaCl, 10% glycerol, 0.05% sarkosyl, 1 mM EDTA, 1 mM DTT). Each wash was incubated for 5 min at 4°C. Bound proteins were eluted with 1.25 CV Strep elution buffer (50 mM HEPES pH 7.6, 250 mM NaCl, 10% glycerol, 0.05% sarkosyl, 1 mM EDTA, 1 mM DTT, 50 mM D-biotin). Peak fractions were identified by SDS-PAGE, pooled, and injected onto an ÄKTA Pure 25 (Cytiva) equipped with a Superdex 75 Increase 10/300 GL (Cytiva) column, operated at a flow rate of 0.5 mL/min using SEC buffer (50 mM Tris pH 7.6, 200 mM NaCl, 10% glycerol, 0.05% sarkosyl, 1 mM EDTA). Peak fractions were identified by SDS-PAGE, aliquoted, and flash frozen in liquid nitrogen before storage at -80°C. Purified proteins were quantified by SYPRO Ruby staining (Invitrogen, S12001) using a standard curve created from acetylated BSA (Promega), according to the manufacturer’s instructions. Final protein concentrations ranged from 0.5 µM for Hotaru FL::His10::TwinStrepII to 2.5 µM for Hotaru ΔMTS::His10::TwinStrepII.

### *In vitro* nuclease activity assays on synthetic oligos

6-carboxyfluorescein (FAM)-labelled DNA substrates were prepared by annealing the oligonucleotides listed in Table 3 and following a previously published protocol^56^. Specifically, 5’-flaps contained FAM-oligo 1 and unlabelled oligos 4 and 2.5, while 3’-flaps contained FAM-oligo 1 and unlabelled oligos 4 and 3.5. Splayed arm substrates were generated from FAM-oligo 1 and unlabelled oligo 4. Nicked Holliday junctions contained FAM-oligo 3 and unlabelled oligos 1.28, 1.32, 2, and 4. Intact Holliday junctions were constructed from FAM-labelled oligo 1 and unlabelled oligos 2, 3, and 4. Double-strand DNA (dsDNA) contained FAM-oligo 1 and unlabelled oligo 1comp. FAM-labelled oligo 5 was used for single-strand DNA (ssDNA). For nuclease activity assays, all protein solutions were prepared on ice as 5X working stocks in enzyme dilution buffer (50 mM Tris pH 7.6 at 4°C, 100 mM NaCl, 10% glycerol, 1 mM EDTA, 1 mM DTT). Nuclease reactions were prepared without protein, incubated at 25°C for 10 min, and then initiated by enzyme addition. Reactions contained 10 nM FAM-labelled DNA substrate and purified proteins (2 nM FL, 2 or 5 nM ΔMTS, or 2 nM Y88F) in cleavage buffer (50 mM Tris-HCl pH 8.0 at 25°C, 2.0 mM MnCl_2_) supplemented with fresh DTT (1 mM) and rAlbumin (0.1 mg/mL) (NEB). Reactions were incubated at 25°C for the indicated times and stopped by incubation with stop buffer (5X = 10 mg/mL proteinase K, 10 mM CaCl_2_, 0.5% SDS) for 30 min at 37°C. Stopped reactions were supplemented with DNA loading dye (6X = 30% glycerol, 0.25% w/v Orange-G) and then electrophoresed through 10% native polyacrylamide gels for 75 min at 150 V. Gels were imaged on a Typhoon FLA 9500 laser-scanning platform using the BPB1 (530DF20) filter (Cytiva). Fluorescent DNA cleavage products were quantified using ImageQuant TL v2005 (Cytiva, GE Healthcare). Product formation is reported as a percentage of total FAM-labelled DNA.

**Table 3.**
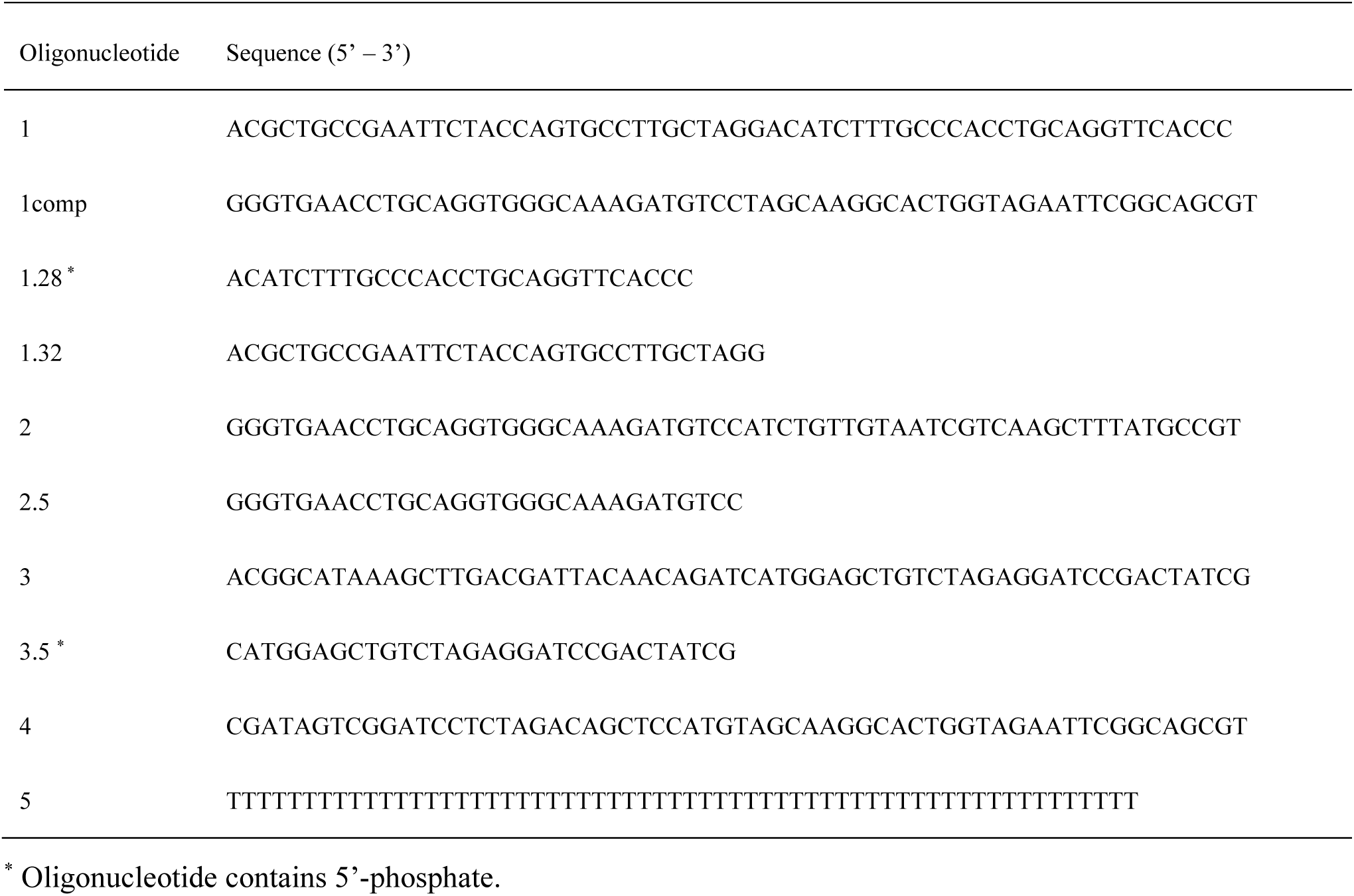
Sequences of the oligonucleotides used to prepare DNA substrates for *in vitro* nuclease assays.

### *In vitro* nuclease activity assays on plasmid

pIRbke8^mut45–47^ was extracted in supercoiled form using QIAfilter Plasmid Midi Kit (Qiagen, 12243), resuspended in cruciform extrusion buffer (50 mM Tris-HCl pH 7.5, 50 mM NaCl, 0.1 mM EDTA), incubated at 37°C for 90 min to induce cruciform extrusion, and stored at 4°C. Immediately before the assay, plasmids were diluted to 10 nM and incubated again at 37°C for 90 min under the same conditions.

All protein solutions were prepared on ice as 5X working stocks in enzyme dilution buffer (50 mM Tris pH 7.6 at 4°C, 100 mM NaCl, 10% glycerol, 1 mM EDTA, 1 mM DTT). Cleavage reactions were prepared without protein, incubated at 25°C for 10 min, and then initiated by enzyme addition. Reactions contained 0.5 nM plasmid DNA and the indicated concentration of purified proteins in cleavage buffer (50 mM Tris-HCl pH 8.0 at 25°C, 2.0 mM MnCl_2_) supplemented with fresh DTT (1 mM) and rAlbumin (0.1 mg/mL) (NEB). Reactions were incubated at 25°C for the indicated times and stopped by incubation with stop buffer (5X = 10 mg/mL proteinase K, 10 mM CaCl_2_, 0.5% SDS) for 30 min at 37°C. Stopped reactions were supplemented with DNA loading dye (6X = 30% glycerol, 0.25% w/v Orange-G) and then electrophoresed through 0.8% agarose gel for 90 min at 90 V at 4°C. Gels were stained with SYBR™ Gold Nucleic Acid Gel Stain (Molecular Probes) and imaged on a Typhoon FLA 9500 laser-scanning platform using the LPB filter (Cytiva). Reaction products were quantified using ImageQuant TL v2005 (Cytiva, GE Healthcare).

EcoRI digestion was used to determine the fraction of plasmid DNA that extruded the cruciform. Reactions containing 0.5 nM of plasmid DNA in 1X rCutSmart^TM^ Buffer (NEB) and 10 U of EcoRI-HF (NEB) were incubated at 37°C for 30 min. Reactions were stopped and analyzed as described above. For the four independent assays performed, the EcoRI-resistant (cruciform-extruding) fraction was 68%, 72%, 73%, and 66%, respectively. Percent species is reported as a percentage of total DNA substrate.

For sequencing the cleaved plasmid by ONT, reactions were incubated for 60 min, stop buffer was skipped, and instead heat-inactivated at 80°C for 20 min. The cleavage product (10 µL) was digested with PstI-HF (NEB, R3140S) at 37°C for 2 h in double reaction volume (20 µL). The downstream sample processing for ONT followed the same protocol as described for mtDNA.

### Protein structure prediction and alignment

The predicted structure of Hotaru was downloaded from the UniProt database (AF-B7YZH2-F1), which hosts AlphaFold-generated structural models^57,58^. The model was evaluated using the confidence metrics provided by AlphaFold, including per-atom confidence, predicted alignment error, and template modeling scores. The highest-confidence model (AF-B7YZH2-F1-v6) was selected for subsequent analyses.

Hotaru is predicted to contain a LEM-3-like GIY-YIG domain spanning residues 85-244 (Pfam PF22945)^34,35^. Structural similarity searches of this region were performed using the DALI server, accessed on December 23, 2025^59^. Specifically, residues 85-244 from the AlphaFold-predicted Hotaru structure were used as the query, and comparisons were limited to entries in the PDB25, with default parameters applied.

### Molecular graphics

Molecular graphics were created with UCSF ChimeraX-1.10^60^, developed by the Resource for Biocomputing, Visualization, and Informatics at the University of California, San Francisco, with support from National Institutes of Health R01-GM129325 and the Office of Cyber Infrastructure and Computational Biology, National Institute of Allergy and Infectious Diseases.

### mtDNA isolation from eggs for long-read ONT sequencing

Unfertilized eggs were collected from apple juice agar plates with yeast paste within 6 h of the previous plate change. Eggs were dechorionated by incubation in 6% bleach for 2 min, rinsed with 0.1% Triton X-100, and collected into DNA LoBind tubes (Eppendorf, 022431021) containing 0.1% Triton X-100. After a brief spin down, 0.1% Triton X-100 was removed and eggs were homogenized with plastic pestles in 100 µL lysis buffer (100 mM Tris-HCl pH 8.0, 10 mM EDTA, 0.5% SDS, 0.25 M NaCl) supplemented with 200 µg/mL proteinase K (NEB, P8107S), followed by incubation at 55 °C for 25 min. Homogenates were mixed with 100 µL 2× SDS-Urea buffer (7 M urea, 0.35 M NaCl, 10 mM Tris-HCl pH 7.8, 1% SDS) and subjected to phenol extraction (pH 8.0; Sigma-Aldrich, P4557) and subsequent chloroform/isoamyl alcohol extraction (24:1; Sigma-Aldrich, 25666). DNA was precipitated by addition of ammonium acetate (0.75 M), glycogen (∼70 µg/mL; Thermo Fisher Scientific, R0561), and 2.5 volumes of ethanol, followed by incubation at -80°C overnight. Precipitated DNA was pelleted, washed with 80% ethanol, air-dried, and resuspended in double-distilled water. DNA was digested with RNase A (Thermo Fisher Scientific, EN0531) and XhoI (NEB, R0146S) at 37°C for 2 h in rCutSmart buffer, and stored at 4°C.

### Long read ONT sequencing

For Oxford Nanopore sequencing, samples were processed as per vendor recommendations using the native barcoding sequencing library prep kit (SQK-NBD114.96) with the exception that the recommended NEBNext FFPE repair step, which is intended to repair nicks, gaps, and other DNA damage, was omitted to avoid it potentially masking Hotaru’s cleavage sites. Briefly, individual samples were treated with NEBNext end-repair module (NEB, E6050), then purified using AMPure beads before undergoing ligation to add barcoding adapters to individual preparations. The resulting barcoded fragments were then pooled and purified before undergoing a second ligation to add the Oxford Nanopore Native Adapter. Sequencing libraries were loaded onto R10.4.1 series (FLO-PRO114M) flowcells and run on a PromethION 24 device for 72 h.

### Mitochondrial reference genome generation

In preliminary analysis, we observed substantial genetic variation between our mitochondrial sequencing data and the mtDNA sequence in the dm6 build of the reference genome. To avoid this genetic variation confounding our analysis, we assembled a new mtDNA reference. This reference sequence was constructed by using minimap2^61^ (2.24-r1122) to align the mtDNA-Y88F sequencing reads to a version of dm6 where chrM was re-oriented to start at the XhoI cut site. Using these alignments, we randomly selected 100 reads that mapped end-to-end to chrM and used these reads to calculate a consensus sequence using abPOA^62^ (v1.5.5). We then replaced chrM in the dm6 reference build with this custom reference to create dm6_mtg_iii, which was used to analyze the Drosophila sequencing experiments. The custom chrM sequence is provided in the software repository that accompanies this work (see **Software availability**).

### Basecalling, trimming, and read mapping

All sequencing runs were base called and demultiplexed with dorado (v1.3.0) using model dna_r10.4.1_e8.2_400bps_hac@v5.2.0 and barcoding kit (SQK-NBD114.96). We observed that doraro’s adapter and barcode trimming algorithm could remove genomic bases from the ends of reads if they have a low-identity alignment to a barcode sequence, which hindered the identification of the precise read start and end points along the genome. We therefore created a custom barcode trimmer (ptrim) designed to only remove barcodes if a sufficiently high identity alignment is found. For the analysis in this paper we required ≥85% identity to remove a barcode or adapter. The trimmed reads were aligned to the corresponding reference sequence for the sample (dm6_mtg_iii for mtDNA samples, pIRbke8 for plasmid samples) using minimap2 with the ‘map-ont’ present. Only read alignments with mapping quality ≥20 were used for subsequent analysis.

### Read end analysis

We reasoned that if Hotaru introduced either a double-strand break or single-strand nick to DNA, it would be apparent in our sequencing data as a population of reads that tended to either initiate (at the 5’ end) or terminate (at the 3’ end) at positions other than the expected XhoI (for mtDNA samples) or PstI (for plasmid samples) site used to linearize the circular molecules. Therefore, we developed a procedure to precisely map the initiation and termination positions for each sequencing read. Using the set of alignments for each read, we transformed the read into a pair of coordinates (*a,b*) where *a* and *b* are the reference positions the first and last base of the read align to, respectively. As potential single-strand nicks may only be apparent on one strand, this coordinate system is stranded. For example, the coordinate (0+, 1000+) indicates a read started at position 0 (the restriction enzyme cut site) on the top reference strand (+) and terminated at position 1000 on the same strand. The coordinate (18000-, 5000-) indicates a read started at position 18,000 on the bottom strand, and terminated at position 5,000 of the same strand. While most reads have a single alignment to the reference genome making this transformation straightforward, some reads had multiple alignments due to sequencing errors, incomplete digestion, or the tendency to form “foldback” reads that switch strands^63^. We therefore considered all alignments that meet our mapping quality threshold when calculating (*a,b*). We require that the read is completely mapped to the reference genome, as truncated alignments (e.g. due to poor sequence quality at read ends) will introduce uncertainty in the initiation or termination position. We therefore discarded any reads where the first or last base was not aligned to the reference, or the set of alignments that did not span the entire read sequence.

### Visualizing DNA cleavage patterns

After transforming the sequencing reads into a set of coordinates, we computed a two-dimensional heatmap depicting the distribution of read start (y-axis) and read end (x-axis) positions along the reference sequence. Each (*a,b*) coordinate was transformed into a binned position and the log10-scaled count within each bin is displayed. For all experiments 100 bins were used in each dimension, corresponding to roughly 188 bp regions of the genome for the mtDNA experiments and 36 bp for the plasmid experiments. The coordinates below the diagonal represent top strand reads (where *a* < *b*), while the coordinates above the diagonal represent bottom strand reads (*b* < *a*).

## Software availability

The complete ONT analysis pipeline and associated code is provided online at https://github.com/jts/hotaru-analysis. The custom barcode trimming software is available at https://github.com/jts/ptrim.

## Data availability

The ONT sequencing data for all experiments have been deposited in the NCBI SRA under BioProject PRJNA1437762.

## Antibodies used in this study

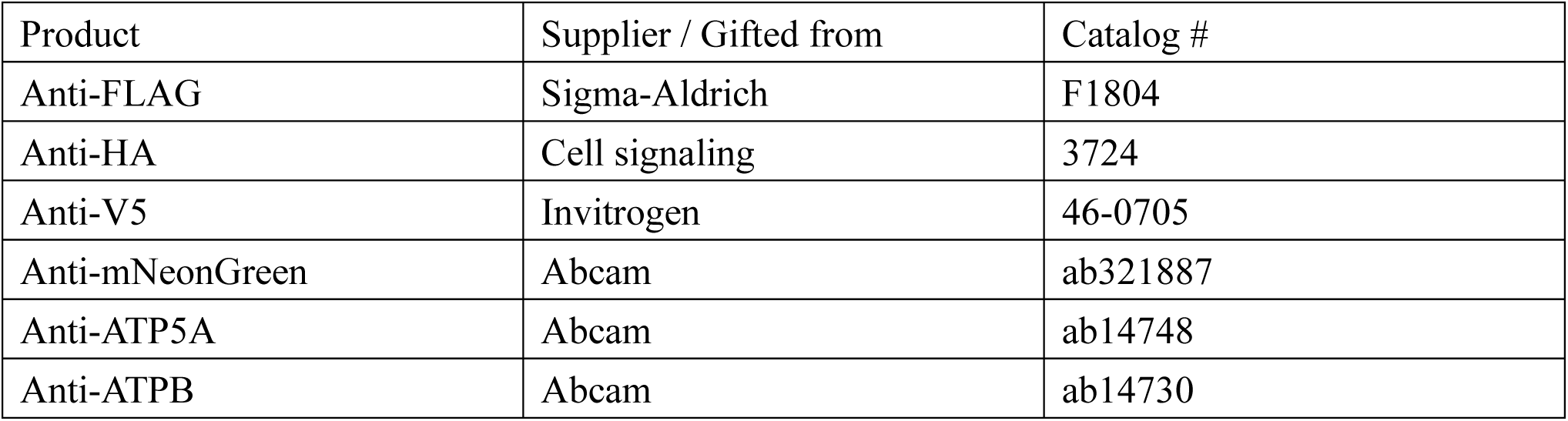
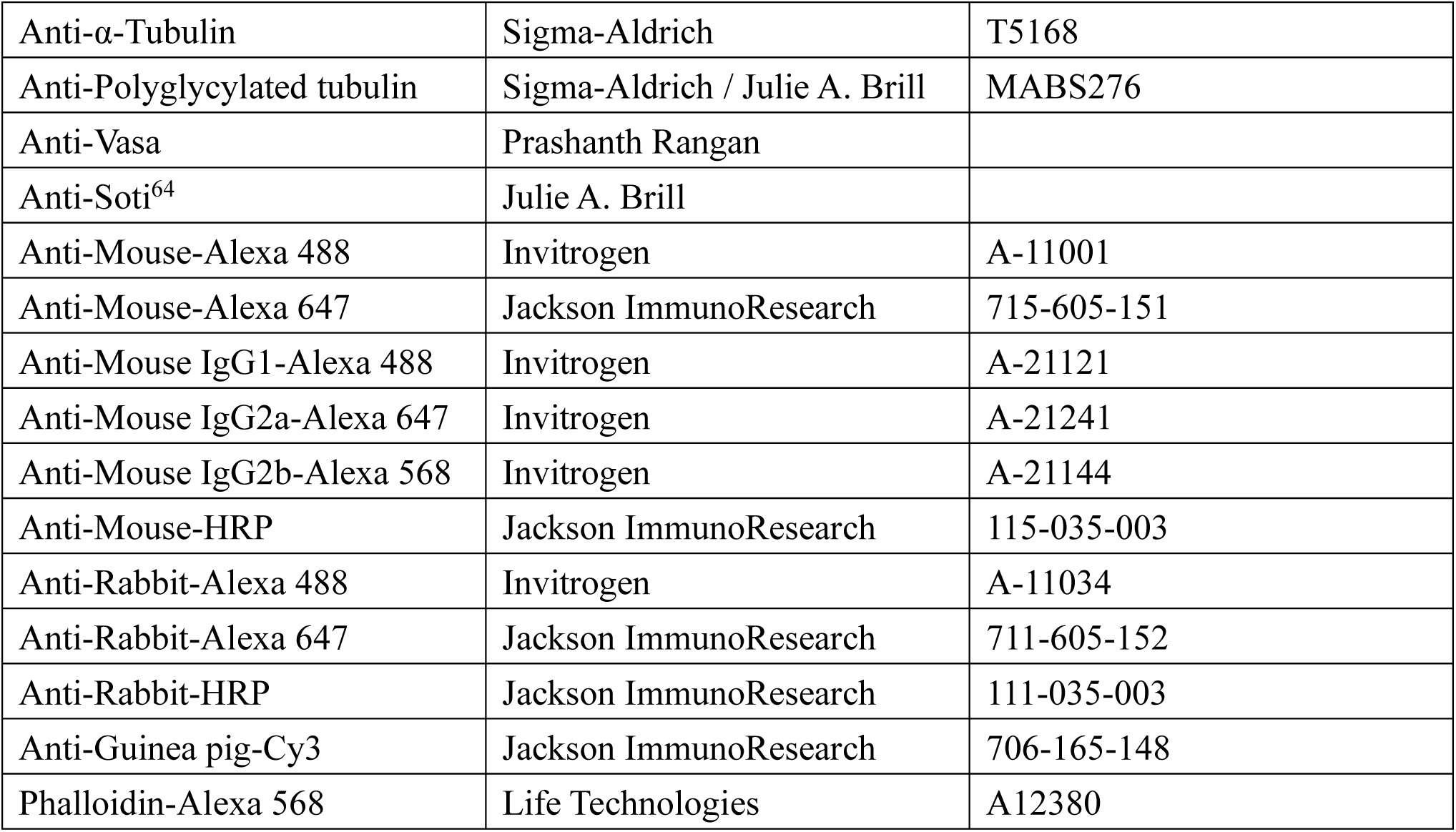

| Product | Supplier / Gifted from | Catalog # |
| --- | --- | --- |
| Anti-FLAG | Sigma-Aldrich | F1804 |
| Anti-HA | Cell signaling | 3724 |
| Anti-V5 | Invitrogen | 46-0705 |
| Anti-mNeonGreen | Abcam | ab321887 |
| Anti-ATP5A | Abcam | ab14748 |
| Anti-ATPB | Abcam | ab14730 |
| Anti-α-Tubulin | Sigma-Aldrich | T5168 |
| Anti-Polyglycylated tubulin | Sigma-Aldrich / Julie A. Brill | MABS276 |
| Anti-Vasa | Prashanth Rangan |  |
| Anti-Soti<sup>64</sup> | Julie A. Brill |  |
| Anti-Mouse-Alexa 488 | Invitrogen | A-11001 |
| Anti-Mouse-Alexa 647 | Jackson ImmunoResearch | 715-605-151 |
| Anti-Mouse IgG1-Alexa 488 | Invitrogen | A-21121 |
| Anti-Mouse IgG2a-Alexa 647 | Invitrogen | A-21241 |
| Anti-Mouse IgG2b-Alexa 568 | Invitrogen | A-21144 |
| Anti-Mouse-HRP | Jackson ImmunoResearch | 115-035-003 |
| Anti-Rabbit-Alexa 488 | Invitrogen | A-11034 |
| Anti-Rabbit-Alexa 647 | Jackson ImmunoResearch | 711-605-152 |
| Anti-Rabbit-HRP | Jackson ImmunoResearch | 111-035-003 |
| Anti-Guinea pig-Cy3 | Jackson ImmunoResearch | 706-165-148 |
| Phalloidin-Alexa 568 | Life Technologies | A12380 |

## Fly stocks used in this study

### Flies obtained from Bloomington Drosophila Stock Center

BL76357, w[1118]; wg[Sp-1]/CyO; MKRS/TM6B, Tb[1]

BL78980, y[1] w[*]; TI{GFP[3xP3.cLa]=CRIMIC.TG4.0}CG11807[CR00674-TG4.0]

CG42391[CR00674-TG4.0-X]/SM6a

BL7879, w[1118]; Df(2R)Exel7135/CyO

BL26072, w[1118]; Mi{GFP[E.3xP3]=ET1}EndoG[MB07150]

BL26551, w[1118]; Df(2R)BSC699, P+PBac{w[+mC]=XP3.RB5}BSC699/SM6a

BL67946, y[1] v[1]; P{y[+t7.7] v[+t1.8]=TRiP.HMS05775}attP2

BL35764, y[1] sc[*] v[1] sev[21]; P{y[+t7.7] v[+t1.8]=TRiP.HMS01512}attP2

BL51852, y[1] v[1]; P{y[+t7.7] v[+t1.8]=TRiP.HMC03426}attP40

BL66533, w[1118]; P{w[+mC]=UAS-mCherry.mito.OMM}3/TM6B, Tb[1]

BL95254, w[*]; mt:ND2[del1]

### Flies gifted from other labs

w^1118^ (Dr. Ruth Lehmann)

;; P{w[+mC]=bam-GAL4::VP16} (Dr. Julie Brill)

;; P{w[+mC]=nos-GAL4::VP16} (Dr. Ruth Lehmann)

;; P{w[+mC]=matα-tub-GAL4::VP16}(Dr. Craig Smibert)

### Flies created in this study

; hotaru^KO^

;; hotaru::mNeonGreen (attP2)

;; hotaru::V5 (attP2)

;; CG42355 UTR-hotaru::V5 (attP2)

;; mst87F UTR-hotaru::V5 (attP2)

;; UASz-hotaru^WT^::3xHA (attP2)

;; UASz-hotaru^Y88F^::3xHA (attP2)

;; UASz-hotaru^E186Q^::3xHA (attP2)

## *hotaru^mut^* allele sequenced from BL78980

ATGTCAAAATTATTAGTATTCAACTTTATCCAAACCATCCCTTTGCAGCTTAGAAATAACTTTAAAAGCAAAGTTAAAGCTAGACCCATTAGTTACTCCCTGGGTACCCTCCACAAGCATCACACAAATTTTTCTACAGAGCTGGAAAATACCATAAATTCTGAGAGCAATTTTAAGAATATCAGTAACCTTATAAAGTTTGTTCGACAACATAGTGGGAGGTTCCGGTGGAAGCGGAGGTAG

## CONTRIBUTIONS

M.S. and T.R.H. conceptualized and designed the study. M.S. performed the experiments unless noted otherwise. H.Y.Y. assisted M.S. with protein purification and performed the *in-vitro* nuclease assays. H.D.M.W. performed computational analysis of the protein structures and assisted M.S. with development of the protein purification protocol. P. C. Z. performed ONT sequencing. J. T. S. analyzed ONT sequencing data. M.S. and T.R.H. wrote the manuscript with input from all authors.

## ACKNOWLEDGEMENTS

We thank Craig A. Smibert, Julie A. Brill, Ruth Lehmann, and the Bloomington Drosophila Stock Center (NIH P40OD018537) for fly stocks; Julie A. Brill and Prashanth Rangan for antibodies; Helen White-Cooper for advice on post-meiotic expression constructs; and Daria E. Siekhaus and Toby Lieber for critical reading and editing of the manuscript. This work was supported by Canadian Institutes of Health Research grant (FRN #542273) to T.R.H. and (FRN #487044) to H.D.M.W. T.R.H. is part of the University of Toronto Medicine by Design initiative, which receives funding from the Canada First Research Excellence Fund. H.D.M.W. is the Tier II Canada Research Chair in Mechanisms of Genome Instability (#CRC-2021-00346). M.S. is supported by the Heiwa Nakajima Foundation Scholarship and the University of Toronto MITO2i Graduate Scholarship. H.Y.Y. is supported by the NSERC Canada Graduate Scholarship. J.T.S. is supported by the Ontario Institute for Cancer Research through funds provided by the Government of Ontario, and the National Human Genome Research Institute (NHGRI project 5R01HG009190).

## REFERENCES

1. Munasinghe, M. & Ågren, J. A. When and why are mitochondria paternally inherited? Current Opinion in Genetics & Development 80, 102053 (2023).

2. Lee, W. et al. Molecular basis for maternal inheritance of human mitochondrial DNA. Nat Genet 55, 1632–1639 (2023).

3. Chen, Z. et al. Mitochondrial DNA removal is essential for sperm development and activity. The EMBO Journal 1–25 (2025) doi:10.1038/s44318-025-00377-5.

4. Nissanka, N., Bacman, S. R., Plastini, M. J. & Moraes, C. T. The mitochondrial DNA polymerase gamma degrades linear DNA fragments precluding the formation of deletions. Nat Commun 9, 2491 (2018).

5. DeLuca, S. Z. & O’Farrell, P. H. Barriers to Male Transmission of Mitochondrial DNA in Sperm Development. Developmental Cell 22, 660–668 (2012).

6. Sagan, L. On the origin of mitosing cells. Journal of Theoretical Biology 14, 225–IN6 (1967).

7. Martijn, J., Vosseberg, J., Guy, L., Offre, P. & Ettema, T. J. G. Deep mitochondrial origin outside the sampled alphaproteobacteria. Nature 557, 101–105 (2018).

8. Mitchell, P. Chemiosmotic coupling in oxidative and photosynthetic phosphorylation. Biol Rev Camb Philos Soc 41, 445–502 (1966).

9. Mitchell, P. Chemiosmotic Coupling in Energy Transduction: A Logical Development of Biochemical Knowledge. in Membrane Structure and Mechanisms of Biological Energy Transduction (ed. Avery, J.) 5–24 (Springer US, Boston, MA, 1973). doi:10.1007/978-1-4684-2016-6_2.

10. Chen, X. J. & Butow, R. A. The organization and inheritance of the mitochondrial genome. Nat Rev Genet 6, 815–825 (2005).

11. Rath, S. P. et al. Mitochondrial genome copy number variation across tissues in mice and humans. Proceedings of the National Academy of Sciences 121, e2402291121 (2024).

12. Wells, J. R. Mitochondrial DNA synthesis during the cell cycle of *Saccharomyces cerevisiae*. Experimental Cell Research 85, 278–286 (1974).

13. Petes, T. D. & Fangman, W. L. Preferential synthesis of yeast mitochondrial DNA in α factor-arrested cells. Biochemical and Biophysical Research Communications 55, 603–609 (1973).

14. Yu, Z., O’Farrell, P. H., Yakubovich, N. & DeLuca, S. Z. The Mitochondrial DNA Polymerase Promotes Elimination of Paternal Mitochondrial Genomes. Current Biology 27, 1033–1039 (2017).

15. Wang, Z. et al. Poldip2 promotes mtDNA elimination during Drosophila spermatogenesis to ensure maternal inheritance. The EMBO Journal 1–25 (2025) doi:10.1038/s44318-025-00378-4.

16. Larsson, N.-G., Oldfors, A., David Garman, J., Barsh, G. S. & Clayton, D. A. Down-Regulation of Mitochondrial Transcription Factor a During Spermatogenesis in Humans. Human Molecular Genetics 6, 185–1991 (1997).

17. Rantanen, A., Jansson, M., Oldfors, A. & Larsson, N.-G. Downregulation of Tfam and mtDNA copy number during mammalian spermatogenesis. Mammalian Genome 12, 787–792 (2001).

18. Zhou, Q. et al. Mitochondrial endonuclease G mediates breakdown of paternal mitochondria upon fertilization. Science 353, 394–399 (2016).

19. Low, R. L. Mitochondrial Endonuclease G function in apoptosis and mtDNA metabolism: a historical perspective. Mitochondrion 2, 225–236 (2003).

20. Ohsato, T. et al. Mammalian mitochondrial endonuclease G. European Journal of Biochemistry 269, 5765–5770 (2002).

21. Strack, P. R. et al. Polymerase delta-interacting protein 38 (PDIP38) modulates the stability and activity of the mitochondrial AAA+ protease CLPXP. Communications Biology 3, 646 (2020).

22. Urrutia, K. et al. DNA sequence and lesion-dependent mitochondrial transcription factor A (TFAM)-DNA-binding modulates DNA repair activities and products. Nucleic Acids Res 52, 14093–14111 (2024).

23. Miller, K. E., Kim, Y., Huh, W.-K. & Park, H.-O. Bimolecular Fluorescence Complementation (BiFC) Analysis: Advances and Recent Applications for Genome-Wide Interaction Studies. Journal of Molecular Biology 427, 2039–2055 (2015).

24. Liu, P. et al. Removal of Oxidative DNA Damage via FEN1-Dependent Long-Patch Base Excision Repair in Human Cell Mitochondria. Mol Cell Biol 28, 4975–4987 (2008).

25. Nishioka, K. et al. Expression and differential intracellular localization of two major forms of human 8-oxoguanine DNA glycosylase encoded by alternatively spliced OGG1 mRNAs. Mol Biol Cell 10, 1637–1652 (1999).

26. Claros, M. G. & Vincens, P. Computational method to predict mitochondrially imported proteins and their targeting sequences. Eur J Biochem 241, 779–786 (1996).

27. Lee, P.-T. et al. A gene-specific T2A-GAL4 library for Drosophila. eLife 7, e35574 (2018).

28. Li, A. Y. Z. et al. Milton assembles large mitochondrial clusters, mitoballs, to sustain spermatogenesis. Proceedings of the National Academy of Sciences 120, e2306073120 (2023).

29. Leader, D. P., Krause, S. A., Pandit, A., Davies, S. A. & Dow, J. A. T. FlyAtlas 2: a new version of the Drosophila melanogaster expression atlas with RNA-Seq, miRNA-Seq and sex-specific data. Nucleic Acids Res 46, D809–D815 (2018).

30. Schäfer, M., Nayernia, K., Engel, W. & Schäfer, U. Translational Control in Spermatogenesis. Developmental Biology 172, 344–352 (1995).

31. Shao, L. et al. Eukaryotic translation initiation factor eIF4E-5 is required for spermiogenesis in Drosophila melanogaster. Development 150, dev200477 (2023).

32. Kuhn, R., Schäfer, U. & Schäfer, M. Cis-acting regions sufficient for spermatocyte-specific transcriptional and spermatid-specific translational control of the Drosophila melanogaster gene mst(3)gl-9. EMBO J 7, 447–454 (1988).

33. Caporilli, S., Yu, Y., Jiang, J. & White-Cooper, H. The RNA export factor, Nxt1, is required for tissue specific transcriptional regulation. PLoS Genet 9, e1003526 (2013).

34. Paysan-Lafosse, T. et al. The Pfam protein families database: embracing AI/ML. Nucleic Acids Res 53, D523–D534 (2025).

35. Blum, M. et al. InterPro: the protein sequence classification resource in 2025. Nucleic Acids Res 53, D444–D456 (2025).

36. Dunin-Horkawicz, S., Feder, M. & Bujnicki, J. M. Phylogenomic analysis of the GIY-YIG nuclease superfamily. BMC Genomics 7, 98 (2006).

37. Song, J., Freeman, A. D. J., Knebel, A., Gartner, A. & Lilley, D. M. J. Human ANKLE1 Is a Nuclease Specific for Branched DNA. Journal of Molecular Biology 432, 5825–5834 (2020).

38. Wyatt, H. D. M., Laister, R. C., Martin, S. R., Arrowsmith, C. H. & West, S. C. The SMX DNA Repair Tri-nuclease. Mol Cell 65, 848–860.e11 (2017).

39. Castor, D. et al. Cooperative control of holliday junction resolution and DNA repair by the SLX1 and MUS81-EME1 nucleases. Mol Cell 52, 221–233 (2013).

40. Wyatt, H. D. M., Sarbajna, S., Matos, J. & West, S. C. Coordinated actions of SLX1-SLX4 and MUS81-EME1 for Holliday junction resolution in human cells. Mol Cell 52, 234–247 (2013).

41. Song, J. et al. Functional dissection of the conserved C. elegans LEM-3/ANKLE1 nuclease reveals a crucial requirement for the LEM-like and GIY-YIG domains for DNA bridge processing. Nucleic Acids Res 53, gkaf265 (2025).

42. Jiang, H. et al. ANKLE1 processes chromatin bridges by cleaving mechanically stressed DNA. Nat Commun 16, 10855 (2025).

43. Payliss, B. J., Patel, A., Sheppard, A. C. & Wyatt, H. D. M. Exploring the Structures and Functions of Macromolecular SLX4-Nuclease Complexes in Genome Stability. Front Genet 12, 784167 (2021).

44. Hurd, T. R. et al. Long Oskar Controls Mitochondrial Inheritance in Drosophila melanogaster. Developmental Cell 39, 560–571 (2016).

45. Lilley, D. M. & Markham, A. F. Dynamics of cruciform extrusion in supercoiled DNA: use of a synthetic inverted repeat to study conformational populations. EMBO J 2, 527–533 (1983).

46. Lilley, D. M. The kinetic properties of cruciform extrusion are determined by DNA base-sequence. Nucleic Acids Res 13, 1443–1465 (1985).

47. Rass, U. et al. Mechanism of Holliday junction resolution by the human GEN1 protein. Genes Dev 24, 1559–1569 (2010).

48. Shimomura, M. & Hurd, T. R. Why and how paternal mitochondrial DNA gets cut out of the inheritance. Current Opinion in Genetics & Development 94, 102381 (2025).

49. Shi, Y. et al. Mammalian transcription factor A is a core component of the mitochondrial transcription machinery. Proc Natl Acad Sci U S A 109, 16510–16515 (2012).

50. Popova, D., Bhide, P., D’Antonio, F., Basnet, P. & Acharya, G. Sperm mitochondrial DNA copy numbers in normal and abnormal semen analysis: A systematic review and meta-analysis. BJOG 129, 1434–1446 (2022).

51. Jenkins, V. K., Larkin, A., Thurmond, J., & FlyBase Consortium. Using FlyBase: A Database of Drosophila Genes and Genetics. Methods Mol Biol 2540, 1–34 (2022).

52. Öztürk-Çolak, A. et al. FlyBase: updates to the Drosophila genes and genomes database. Genetics 227, iyad211 (2024).

53. DeLuca, S. Z. & Spradling, A. C. Efficient Expression of Genes in the Drosophila Germline Using a UAS Promoter Free of Interference by Hsp70 piRNAs. Genetics 209, 381–387 (2018).

54. Shao, Z. et al. Spatially revealed roles for lncRNAs in Drosophila spermatogenesis, Y chromosome function and evolution. Nat Commun 15, 3806 (2024).

55. Fitzgerald, D. J. et al. Protein complex expression by using multigene baculoviral vectors. Nat Methods 3, 1021–1032 (2006).

56. Tse, Y. W. E., Yun, H. Y. & Wyatt, H. D. M. Annealing and purification of fluorescently labeled DNA substrates for in vitro assays. STAR Protoc 4, 102128 (2023).

57. Fleming, J. et al. AlphaFold Protein Structure Database and 3D-Beacons: New Data and Capabilities. J Mol Biol 437, 168967 (2025).

58. Jumper, J. et al. Highly accurate protein structure prediction with AlphaFold. Nature 596, 583–589 (2021).

59. Holm, L. Dali server: structural unification of protein families. Nucleic Acids Res 50, W210–W215 (2022).

60. Meng, E. C. et al. UCSF ChimeraX: Tools for structure building and analysis. Protein Sci 32, e4792 (2023).

61. Li, H. Minimap2: pairwise alignment for nucleotide sequences. Bioinformatics 34, 3094–3100 (2018).

62. Gao, Y. et al. abPOA: an SIMD-based C library for fast partial order alignment using adaptive band. Bioinformatics 37, 2209–2211 (2021).

63. Heinz, J. M., Meyerson, M. & Li, H. Detecting foldback artifacts in long-reads. BMC Genomics 27, 144 (2026).

64. Kaplan, Y., Gibbs-Bar, L., Kalifa, Y., Feinstein-Rotkopf, Y. & Arama, E. Gradients of a ubiquitin E3 ligase inhibitor and a caspase inhibitor determine differentiation or death in spermatids. Dev Cell 19, 160–173 (2010).

